# Regulation of intracellular cAMP levels in osteocytes by mechano-sensitive focal adhesion kinase via PDE8A

**DOI:** 10.1101/2024.06.28.601153

**Authors:** Garyfallia Papaioannou, Tadatoshi Sato, Caroline Houghton, Parthena E Kotsalidis, Katelyn E Strauss, Thomas Dean, Alissa J. Nelson, Matthew Stokes, Thomas J Gardella, Marc N Wein

## Abstract

Osteocytes are the primary mechano-sensitive cell type in bone. Mechanical loading is sensed across the dendritic projections of osteocytes leading to transient reductions in focal adhesion kinase (FAK) activity. Knowledge regarding the signaling pathways downstream of FAK in osteocytes is incomplete. We performed tyrosine-focused phospho-proteomic profiling in osteocyte-like Ocy454 cells to identify FAK substrates. Gsα, parathyroid hormone receptor (PTH1R), and phosphodiesterase 8A (PDE8A), all proteins associated with cAMP signaling, were found as potential FAK targets based on their reduced tyrosine phosphorylation in both FAK-deficient or FAK inhibitor treated cells. Real time monitoring of intracellular cAMP levels revealed that FAK pharmacologic inhibition or gene deletion increased basal and GPCR ligand-stimulated cAMP levels and downstream phosphorylation of protein kinase A substrates. Mutating FAK phospho-acceptor sites in Gsα and PTH1R had no effect on PTH- or FAK inhibitor-stimulated cAMP levels. Since FAK inhibitor treatment augmented cAMP levels even in the presence of forskolin, we focused on potential FAK substrates downstream of cAMP generation. Indeed, PDE8A inhibition mimicked FAK inhibition at the level of increased cAMP, PKA activity, and expression of cAMP-regulated target genes. *In vitro* kinase assay showed that PDE8A is directly phosphorylated by FAK while immunoprecipitation assays revealed intracellular association between FAK and PDE8A. Thus, FAK inhibition in osteocytes acts synergistically with signals that activate adenylate cyclase to increase intracellular cAMP. Mechanically-regulated FAK can modulate intracellular cAMP levels via effects on PDE8A. These data suggest a novel signal transduction mechanism that mediates crosstalk between mechanical and cAMP-linked hormonal signaling in osteocytes.

## Introduction

Mimicking the skeletal benefits of weight-bearing exercise through molecular signals is a major goal in bone biology. Exercise boosts bone formation and bone mass [1] while immobilization or lack of gravity, through a reduction of mechanical loading, lead to a reduction of bone mass [2]. Osteocytes play a central role in orchestrating how bone senses mechanical and hormonal signals [3, 4]. Osteocytes detect mechanical loading as changes in fluid flow shear stress (FFSS) across their dendritic projections [5]. FFSS or mechanical loading stimulates bone anabolism by downregulating sclerostin, an osteoblast inhibitor [6].

Hormonal cues also reduce sclerostin expression by osteocytes. Parathyroid hormone (PTH), a bone anabolic agent when delivered via once daily subcutaneous injections [7], binds to its Gsα- linked G-protein-coupled-receptor (PTH1R) in osteocytes, increases cAMP levels and inhibits sclerostin expression [3]. Therefore, both GPCR signaling, and mechanical cues converge to regulate sclerostin expression. Furthermore, through multiple proposed mechanisms [8–10], mechanical disuse impairs the bone anabolic effects of intermittent PTH injections, a commonly used osteoporosis treatment. On the contrary, the combination of exercise and PTH may maximize the effects of either treatment alone [11]. For example, combined intermittent PTH treatment with whole body vibration led to enhanced gains in spine bone density compared to PTH alone [12]. Despite these observations, little is known about how mechanical and hormonal signals are integrated within osteocytes. Molecular mechanisms that mimic mechanical signals could be used as drug targets to increase bone mass and enhance the effects of PTH in bone anabolism.

We previously described a key role of the tyrosine kinase Focal Adhesion Kinase (FAK) in the intracellular signaling pathway downstream of FFSS [13]. FAK is an integral part of focal adhesions, complex structures that assemble at the plasma membrane where integrins interact with the extracellular matrix (ECM) [14]. FAK is activated downstream of integrin/ECM interaction [15]. Relative to other cells, osteocytes show constitutively-high FAK signals due to ‘outside-in’ signaling from their surrounding bone matrix via integrins [13]. FFSS rapidly causes integrin-extracellular matrix dissociation, thereby reducing FAK activity. When active, FAK phosphorylates class IIa histone deacetylase proteins (HDAC4 and HDAC5) and promotes their cytoplasmic retention. When FAK kinase activity is suppressed transiently by FFSS, HDAC4/HDAC5 tyrosine phosphorylation is reduced, leading to their subsequent nuclear translocation. In the nucleus, class IIa HDACs suppress MEF2-driven expression of sclerostin (encoded by the SOST gene) [13]. Since sclerostin normally suppresses bone formation by osteoblasts [6], FAK inhibition could lead to increased bone formation by reducing sclerostin levels. Notably, a dual FAK/PYK2 inhibitor increases bone anabolism and bone mass in female rats with bone loss due to surgical hypogonadism [16]. Despite these advances, further development of FAK inhibitors to treat osteoporosis is limited by lack of knowledge regarding the full suite of FAK substrates in osteocytes.

In this study, we demonstrate that FAK regulates signaling downstream of Gsa-coupled GPCRs including PTH1R. Tyrosine-focused phospho-proteomic profiling demonstrated that several proteins involved in cAMP signaling showed reduced phosphorylation in response to FAK inhibition or gene deletion. Cells lacking FAK or with inhibited FAK activity have elevated basal and ligand-stimulated cAMP levels and increased signaling downstream of cAMP. Phosphodiesterase 8A (PDE8A) is a direct FAK substrate that controls cAMP levels in bone cells. Therefore, mechanically-regulated FAK inhibition may play a role in increasing the bone anabolic effects of hormonal signaling. Our data suggest a novel mechanism of crosstalk between mechanical and cAMP-linked hormonal signaling in osteocytes.

## Results

### FAK controls phosphorylation of proteins associated with cAMP signaling

To understand how FAK regulates bone anabolism, we sought to identify novel targets of this tyrosine kinase in osteocytes. Phospho-tyrosine-focused phosphoproteomic profiling was performed to identify substrates of FAK in an osteocyte-like cell line (Ocy454 cells). Tryptic peptides were isolated from cells treated ± the FAK inhibitor VS6063 [17] or ± FAK gene deletion (**Figure 1a,b**). 432 proteins showed reduced tyrosine phosphorylation in FAK KO and FAK inhibitor-treated cells compared to control (**Figure 1c, Supplemental Table 1**). FAK itself and paxillin are known FAK substrates; thus, our ability to detect reduced phosphorylation of these proteins validates this approach. Among proteins whose tyrosine phosphorylation was reduced by both VS6063 and FAK deletion, Gsα and PTH1R (**Supplemental Figure 1**) were of particular interest due to their known roles in bone biology [18–20]. In addition, other proteins related to cAMP signaling were identified, including phosphodiesterase 8A (PDE8A) (**Figure 1d**).

**Figure 1:**
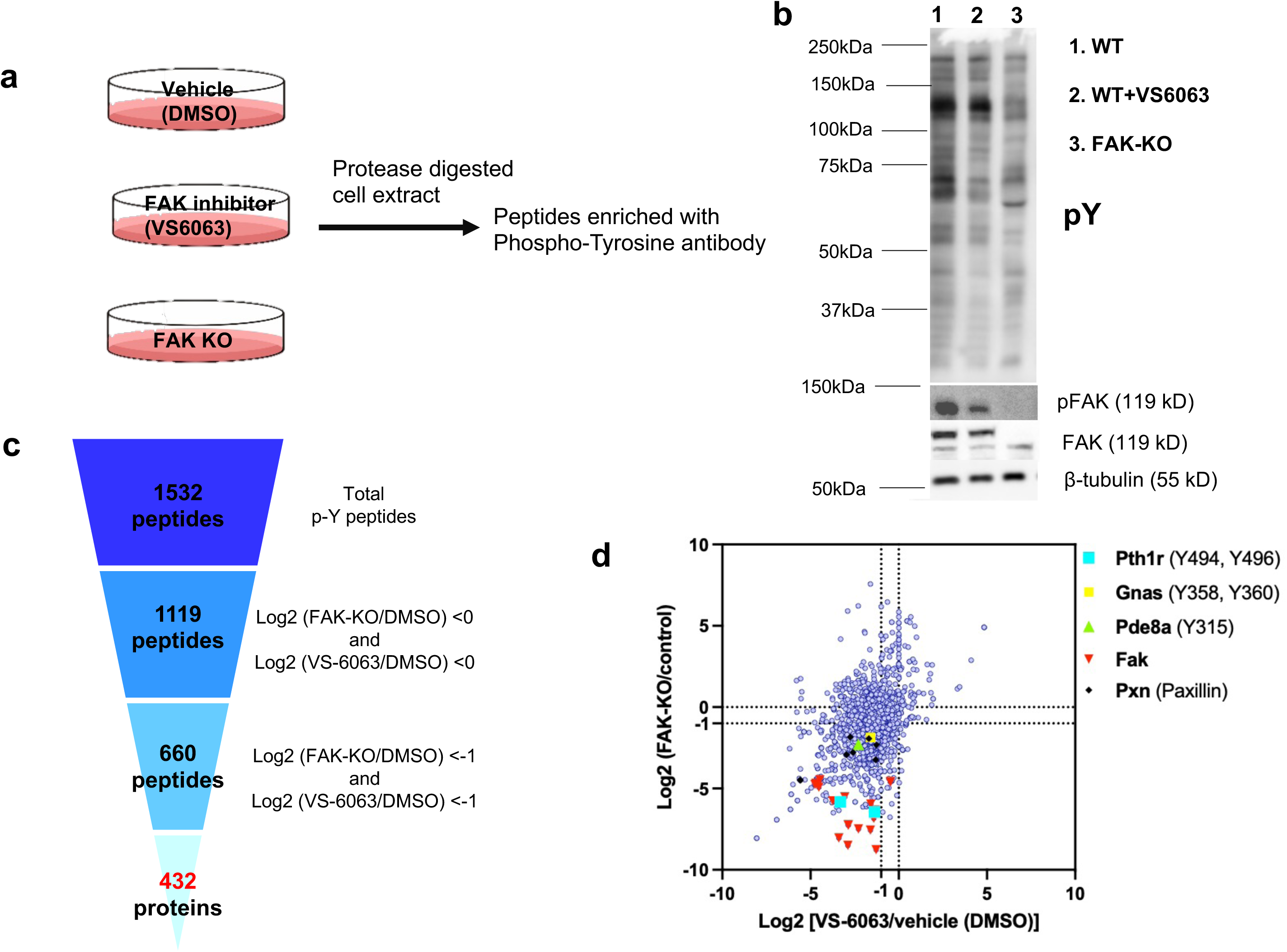
Identification of Focal Adhesion Kinase substrates in Ocy454 cells. (**a**) Experimental design of phosphoproteomic profiling in Ocy454 cells. (**b**) Immunoblotting with antibodies for phosphorylated tyrosine residues, pFAK, total FAK and ß-tubulin. Protein lysates were isolated from control, FAK inhibitor (VS6063, 10 µM, 30 minutes) treated, and FAK KO Ocy454 cells. (**c**) 432 proteins showed reduced abundance of tyrosine phosphorylated residues in FAK KO and FAK inhibitor-treated cells compared to the control. (**d**) Dot plot showing the abundance of tyrosine phosphorylation sites in FAK KO cells and FAK inhibitor treated cells versus control. Left lower quadrant: proteins with reduced abundance of tyrosine phosphorylated residues in both comparisons. The phosphorylation sites of interest are shown in parenthesis. FAK and Pxn (known Fak targets) are positive controls.

### FAK inhibition increases basal and ligand-stimulated cAMP levels

Having identified cAMP signaling related proteins regulated by FAK, we asked if FAK could control Gsα signaling. We first used HEK293 cells harboring a GloSensor system [21] which generates luminescence proportionate to the amount of intracellular cAMP. In this system, VS6063 treatment rapidly increased intracellular cAMP levels in a dose-dependent manner in the absence of exogenously-added GPCR ligand (**Figure 2a**). Furthermore, VS6063 pre-treatment augmented agonist (isoproterenol, PGE_2_, or PTH)-stimulated cAMP levels (**Figure 2b-c**). HEK293 cells express endogenous receptors for isoproterenol and PGE_2_. To detect PTH-induced cAMP increases in these cells, we used cells stably expressing PTH1R [22, 23]. Similar findings were observed using four distinct pharmacologic FAK inhibitors (PF-431396, PF-562271, GSK2256098, Y15) (**Figure 2c**).

**Figure 2:**
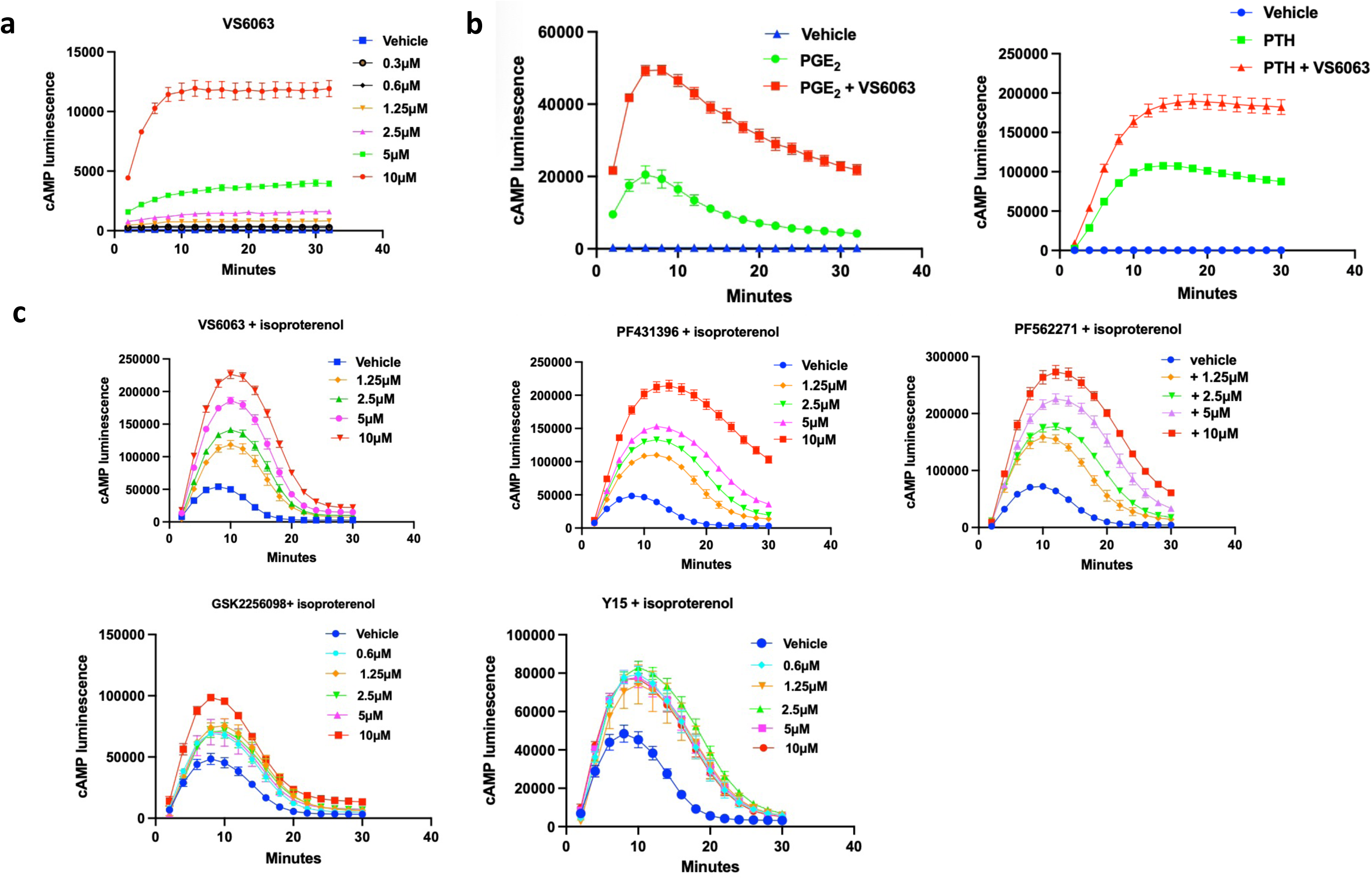
FAK inhibitors increase cAMP levels in HEK293 cells at baseline and upon GPCR- ligand stimulation. GloSensor cAMP luminescence over time is shown (**a**) Dose response of VS6063 on cAMP in GloSensor-expressing HEK cells. (**b**) GloSensor-expressing HEK293 cells treated with PGE2 (10^-3^ M) or PTH (200 pM, a sub-maximal dose) with or without FAK inhibitor (10 μM). GloSensor HEK293 cells stably expressing PTH1R were used for the PTH treatment. (**c**) Dose response of different FAK inhibitors in GloSensor-expressing HEK cells treated with isoproterenol [10^-6^ M].

To extend the relevance of these findings to bone cells, we used GloSensor-expressing osteoblast-like Saos2 cells which express endogenous PTH receptors [24]. Similar to HEK293 cells, VS6063 treatment increased cAMP levels in Saos2 cells (**Figure 3a**). Cilengitide is an RGD peptide [25] functioning upstream of outside-in integrin signaling which activates FAK. Disrupting ECM/integrins interactions using cilengitide reduces FAK activity in Saos2 cells and in osteocyte-like cells [13]. As predicted by our model, cilengitide-induced FAK suppression increased cAMP levels similarly to FAK inhibitors, confirming the effect of the integrin/FAK signaling pathway on cAMP (**Figure 3a**). Similar to results obtained in PTH1R-expressing HEK293 cells, VS6063 treatment potentiated PTH-induced increases in cAMP levels in Saos2 cells (**Figure 3b**). Ocy454 cells are a conditionally-immortalized osteocyte-like cell line [26]. Ocy454 cells were transiently transfected with the GloSensor cDNA and then treated with isoproterenol ± VS6063. Similar to results in HEK293 and Saos2 cells, FAK inhibition potentiated ligand-induced cAMP increases in Ocy454 cells (**Figure 3c**).

**Figure 3:**
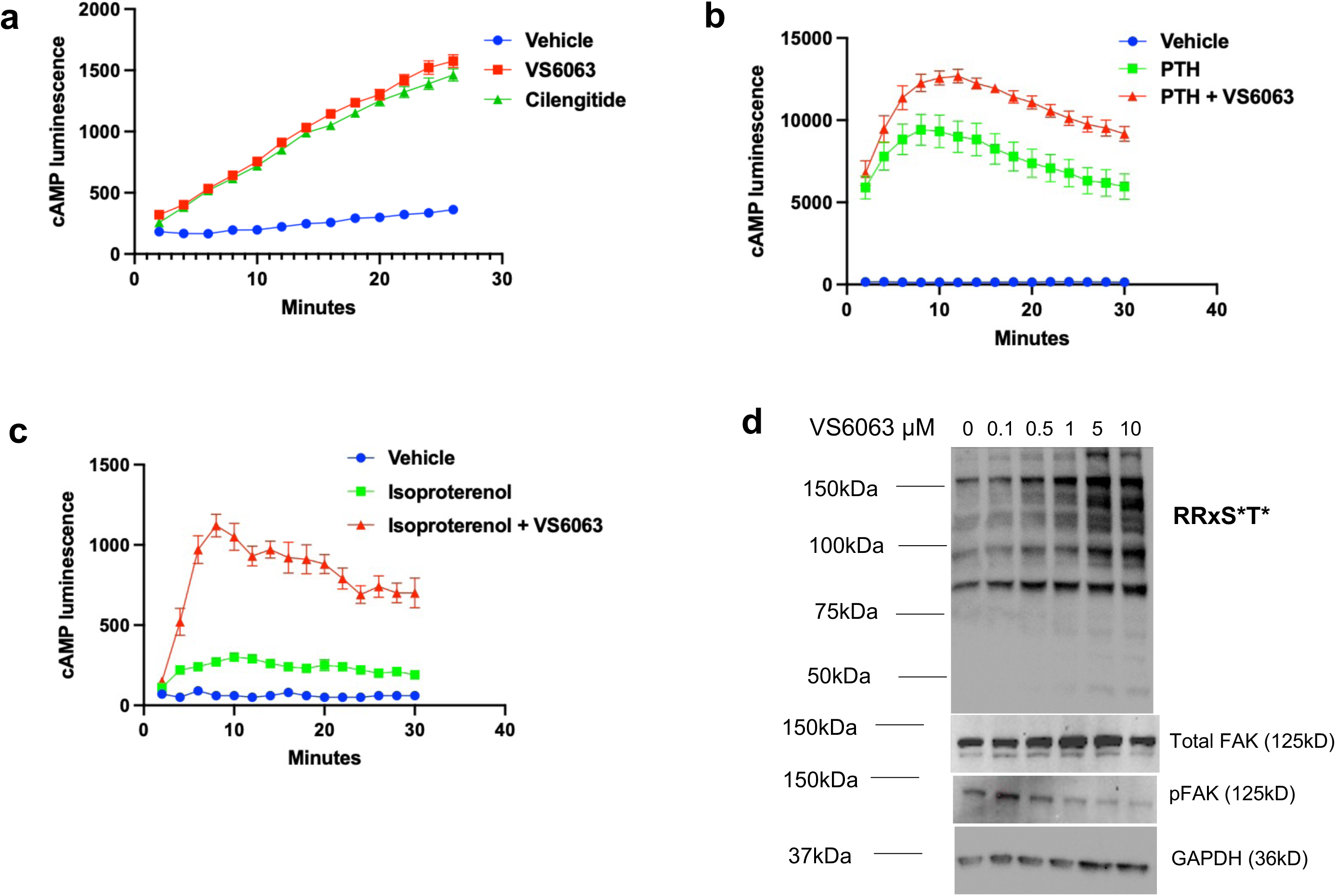
FAK inhibition increases cAMP levels in osteoblast- and osteocyte-like cells at baseline and upon PTH stimulation. (**a**) Saos2 cells treated with FAK inhibitor VS6063 (10 µM) or cilengitide (100 μM). (**b**) Saos2 cells treated with PTH with or without FAK inhibitor VS6063 (10 μM). (**c**) Ocy454 cells transiently expressing GloSensor treated with isoproterenol with or without FAK inhibitor VS6063 (10 μM). cAMP luminescence over time is shown in GloSensor assay. Vehicle is DMSO. (**d**) Immunoblotting for phosphorylated PKA substrates in Ocy454 cells treated with VS6063 (dose response). Cells were treated for 30min.

Protein Kinase A (PKA) is a major downstream effector of cAMP [27]. The effect of FAK inhibition on PKA activation was demonstrated by measuring the signal of phosphorylated PKA (pPKA) substrates (using a degenerate antibody which detects all proteins bearing the RRxS*/T* PKA consensus phosphorylation motif) in Ocy454 cells treated with FAK inhibitor. This orthologous approach demonstrated increased pPKA substrate signal following VS6063 treatment (**Figure 3d)**.

To confirm findings seen with pharmacologic FAK inhibition, we next used genetic approaches to delete or reduce FAK expression. First, FAK deficient GloSensor HEK293 cells were generated by CRISPR/Cas9-mediated gene deletion (**Supplemental Figure 2**). As predicted by our studies with VS6063, FAK-deficient cells showed increased isoproterenol-stimulated cAMP levels compared to control cells (**Figure 4a**). In addition, FAK knockdown GloSensor HEK293 cells were generated via lentiviral-mediated shRNA knockdown. Similar to the effects of CRISPR-mediated FAK gene deletion, shRNA-induced FAK knockdown led to increased isoproterenol-stimulated cAMP levels compared to cells infected with lentivirus containing empty vector (**Figure 4b**). Therefore, genetic and pharmacologic loss of function data indicates that FAK inhibition increases cAMP levels both at baseline and in response to Gsα-coupled GPCR agonists.

**Figure 4:**
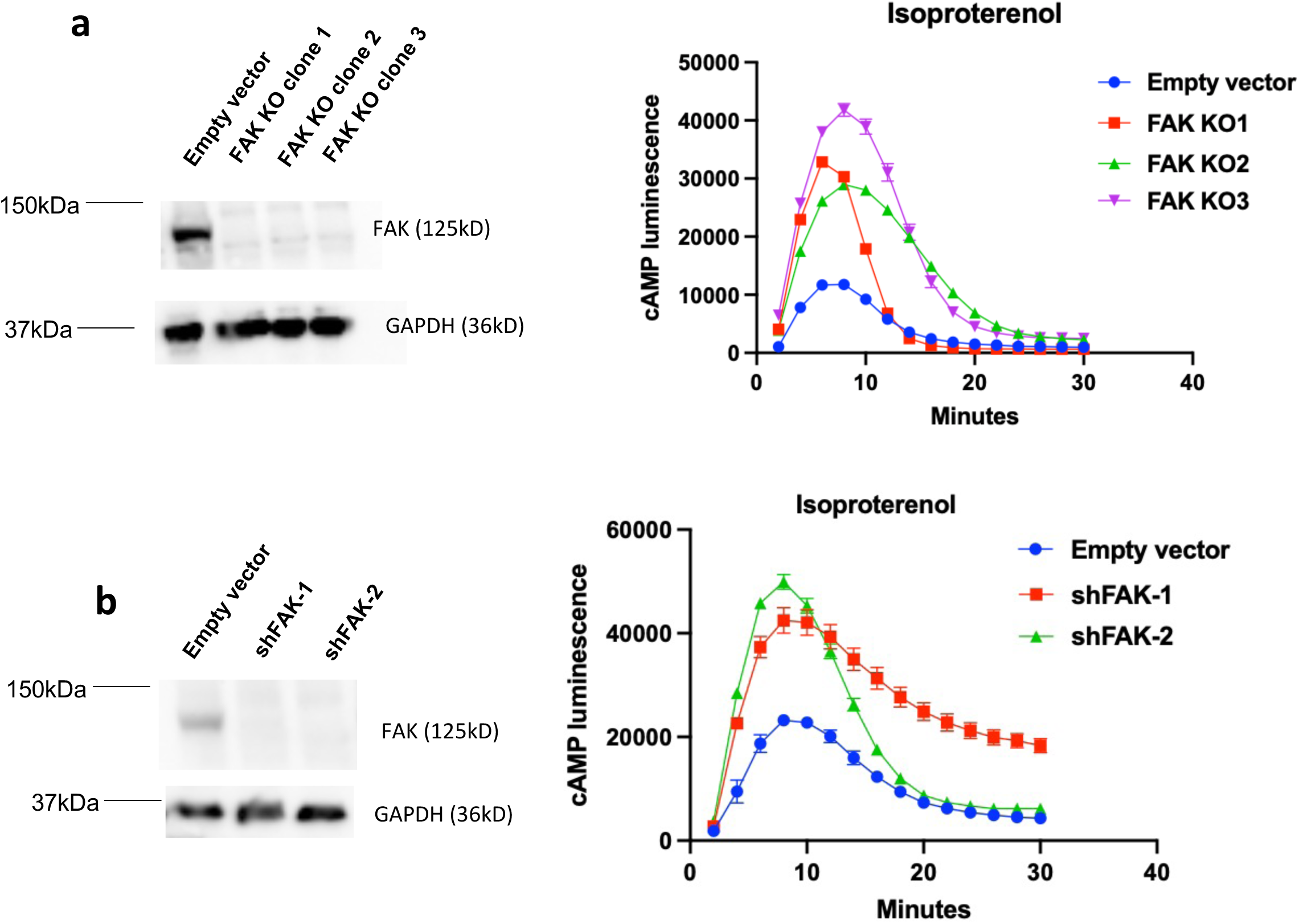
FAK genetic perturbation increases cAMP levels in HEK293 cells upon stimulation with GPCR ligands. (**a**) <u>Left</u>: Immunoblotting for FAK in protein lysates from FAK KO cells generated via CRISPR/Cas9 (clones 1-3). <u>Right</u>: GloSensor assay revealed increased cAMP luminescence in FAK KO cells compared to control cells upon treatment with isoproterenol (10^-6^ M). Control cells were transfected with the empty vector PX458. (**b**) <u>Left</u>: Immunoblotting for FAK in protein lysates from cells transduced with the indicated lentiviruses. <u>Right</u>: GloSensor assay revealed increased cAMP luminescence in FAK knockdown cells (shRNAs 1-2) compared to control cells upon treatment with isoproterenol (10^-6^ M). Control cells were infected with the empty pLKO.1 vector.

### FAK regulates cAMP signaling downstream of Gsα

Next, we sought to understand the mechanism through which FAK controls cAMP levels. First, we focused on Gsα itself since pharmacologic and genetic FAK inhibition augments cAMP levels at ‘baseline’ and in response to several Gsα-coupled receptors. FAK-regulated phosphorylation of two Gsα tyrosine residues (Y358 and Y360) was detected in our phospho-proteomic profiling study. Notably, these residues are located near the Gsα/GPCR interface in the model shown in **Figure 5a** [28] thus their regulated tyrosine phosphorylation may affect GPCR/G protein coupling. However, a tyrosine-to-phenylalanine mutated Gsα (2YF) construct introduced into Gsα deficient HEK293 cells showed similar effects on basal and ligand-stimulated cAMP levels as a wild type Gsα expression construct (**Figure 5b**).

**Figure 5.**
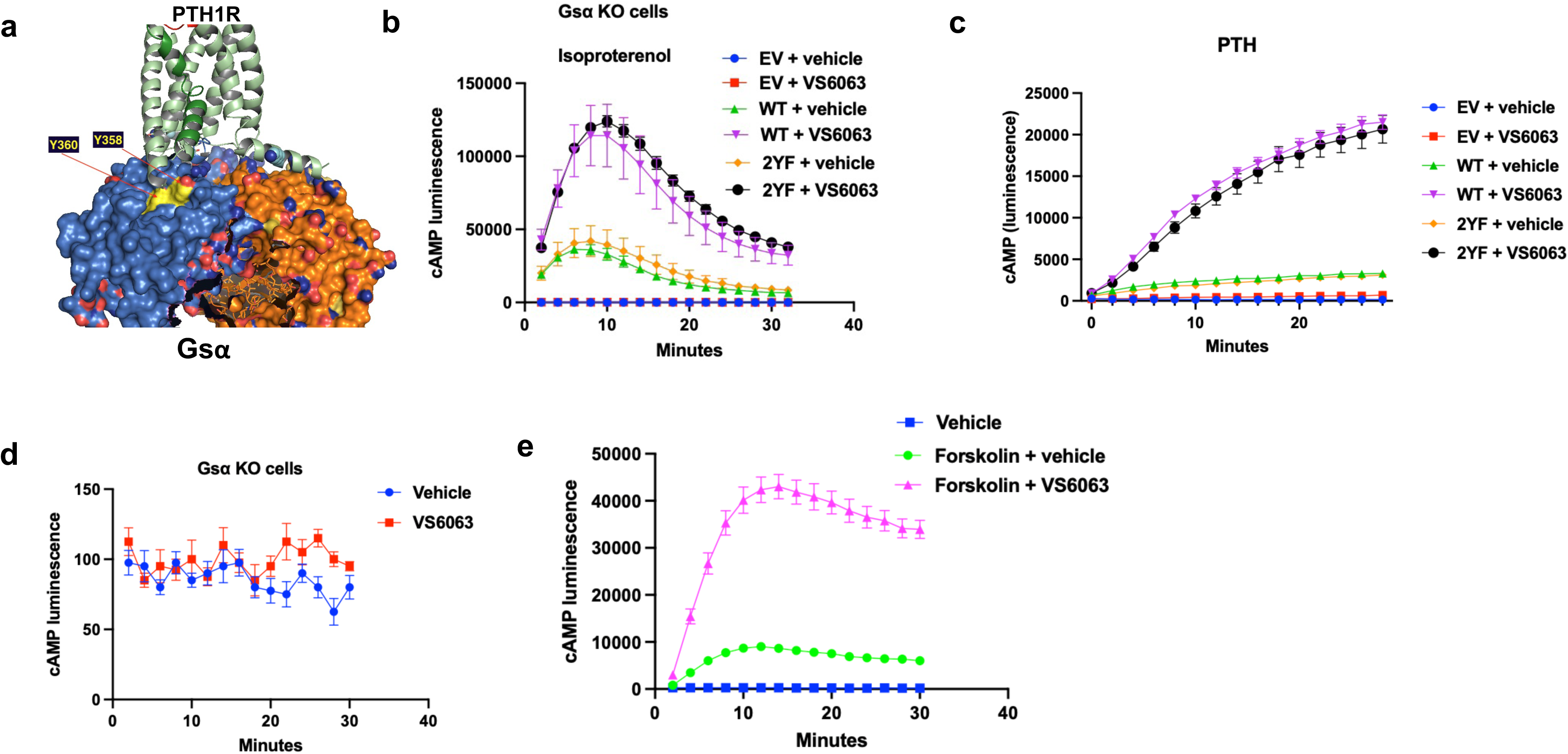
The regulation of cAMP by FAK likely occurs via a mechanism downstream of adenyl-cyclase: (**a**) PTH1R-Gsα/ß/γ cryo structure. Y358 and Y360 residues of Gsα are located at the interface with the IC-3 loop of the receptor and may be contacting residues in that loop or the flanking TM5 and TM6. The receptor is green with TM6 darker green and kinked in the middle for the active state. Gsα is blue and ß/γ are orange. Y358 and Y360 of Gsα are yellow for carbons and red for Oxygens. The sidechain OH of 358 is exposed to the IC3 loop from the outside while the OH of 360 is accessed best from the other side of the complex and can be seen in the 180° rotated view on the right. This model that was generated in PyMOL software using PDB datafile 6NBF. (**b**) cAMP luminescence over time in Gsα KO cells transfected with wild type (WT) or 2YF mutant (Y358F/Y360F) Gsα plasmids with or without FAK inhibitor. (**c**) cAMP luminescence over time in HEK293 cells transfected with WT or 2YF mutant (Y494F/Y496F) PTH1R plasmids ± FAK inhibitor (10 μM). (**d**) cAMP luminescence over time in Gsα KO cells treated with vehicle or FAK inhibitor (10 μM). (**e**) cAMP luminescence over time in HEK293 cells treated with forskolin (10 μM) ± FAKi (10 μM).

We next interrogated the effects of PTH1R C-terminal tail tyrosine phosphorylation (Y494 and Y496) sites using a similar mutagenesis strategy. PTH1R is phosphorylated on its cytoplasmic tail at Y494 by insulin-like growth factor type I receptor, a modification that may regulate PTH1R localization and function [29]. Although PTH1R tyrosine phosphorylation likely does not explain changes in basal or isoproterenol/PGE_2_-stimulated cAMP levels, these studies were performed based on the central role of PTH in bone metabolism and osteocyte biology [30]. Similar to Gsα, tyrosine to phenylalanine mutagenesis of PTH1R did not affect PTH-induced cAMP induction (**Figure 5c**). Furthermore, since GPCR C-terminal tails play important roles in ß-arrestin recruitment and receptor internalization [31], we tested this 2YF PTH1R construct in PTH1R internalization and ß-arrestin recruitment and saw no obvious differences between the mutated and the wild type receptor (**Supplemental Figure 3a)**. In addition, FAK inhibitors did not cause changes in PTH1R internalization (**Supplemental Figure 3b**) or Gαq/PKC-dependent increases in intracellular calcium (**Supplemental Figure 3c**). Therefore, FAK-mediated changes in cAMP levels are unlikely to occur via direct effects on Gsα and PTH1R at the specific tyrosine sites detected in our phosphoproteomic dataset. At present, we cannot rule out lack of sensitivity to detect effects of point mutations due to artifacts associated with these overexpression systems.

To pinpoint the level(s) in the cAMP signaling cascade at which FAK inhibitors exert their effects, we treated Gsα KO cells with VS6063 and, unlike the case in control cells (**Figure 2a**), did not observe increases in cAMP levels (**Figure 5d**). Therefore, it is unlikely that FAK inhibition directly activates adenylate cyclase in the absence of Gsα. In contrast, VS6063 treatment boosted cAMP levels generated by forskolin, a direct adenylate cyclase activator (**Figure 5e**). Notably, VS6063 was tested *in vitro* against a panel of recombinant phosphodiesterases and did not significantly inhibit PDE assay at doses similar to our cell culture studies (**Supplemental Table 2**), thus making it unlikely that ‘off-target’ direct PDE inhibition by VS6063 is responsible for the cAMP-increasing effects of this compound. Taken together, these findings suggest that the molecular action of FAK in cAMP signaling occurs *downstream* of adenylate cyclase.

### Phosphodiesterase 8A is a FAK target that regulates SOST expression in osteocytes

We next turned our attention to phosphodiesterase 8A (PDE8A), a selective cAMP phosphodiesterase [32] that was also identified as a possible FAK substrate in our phosphoproteomic study at position Y315 (**Figure 1d**). GloSensor Saos2 cells were treated with the selective PDE8A inhibitor PF04957325 [33] ± isoproterenol or PTH. Both basal and ligand-stimulated cAMP levels were increased by PDE8A inhibitor treatment (**Figure 6a**). To confirm these pharmacologic findings, PDE8A-deficient GloSensor HEK293 cells were generated. These cells demonstrated increased isoproterenol-stimulated cAMP levels compared to control cells (**Figure 6b**). Moreover, PDE8A inhibitor treatment led to reduced SOST expression in Ocy454 (**Figure 6c**) in a manner like that of FAK inhibitor treatment [13].

**Figure 6:**
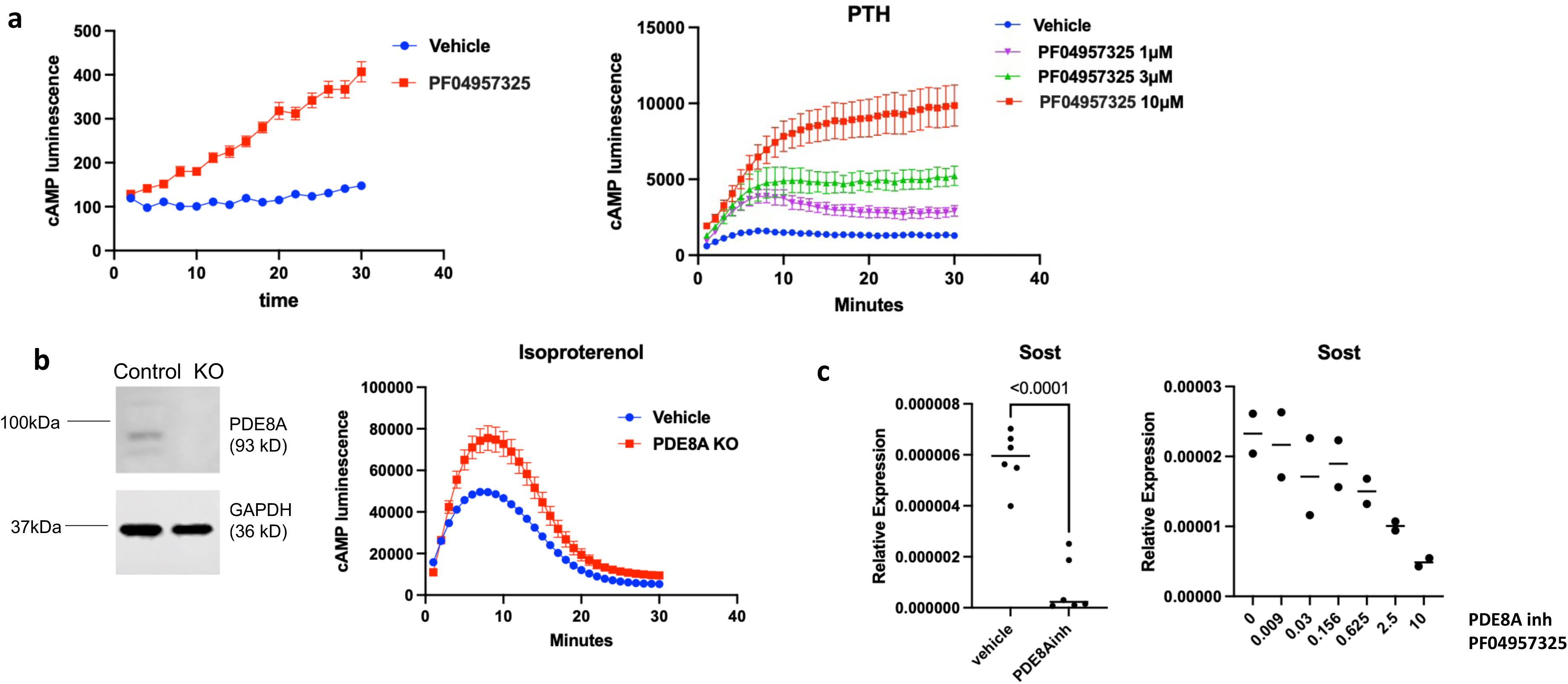
PDE8A inhibition regulates cAMP levels similarly to FAK inhibition: (**a**) <u>Left:</u> GloSensor cAMP assay in Saos2 cells following PDE8A inhibitor treatment (PF04957325, 10 μM). <u>Right:</u> Dose response of the PDE8A inhibitor PF04957325 in cAMP in GloSensor- expressing Saos2 cells. cAMP luminescence over time is shown. (**b**) <u>Left:</u> PDE8A immunoblotting in PDE8A KO cells. <u>Right</u>: GloSensor assay revealed increased cAMP luminescence in PDE8A KO cells compared to control cells upon treatment with isoproterenol (10^-6^ M). Control cells were transfected with the empty vector PX459. (**c**) qRT-PCR showing relative expression of *Sost* ± treatment with PDE8A inhibitor 10 μM (left) and dose response of PDE8A inhibitor (right). Cells were treated for 4 hours prior to RNA isolation. For each group in control vs PDE8A inhibitor treated cells n = 6 biologic replicates for RNA and two-sided unpaired t test were used. N=2 biologic replicates were used for the dose response experiment.

Next, we studied the molecular mechanism linking FAK and PDE8A. In an *in vitro* kinase assay using recombinant FAK and PDE8A proteins, we observed FAK autophosphorylation and FAK-dependent PDE8A tyrosine phosphorylation (**Figure 7a**). Subsequently, co-immunoprecipitation studies demonstrated that FAK and PDE8A associate in cells (**Figure 7b-c**). Interestingly, in a cell-free system we did not detect changes in PDE8A phosphodiesterase activity following its phosphorylation by FAK (**Supplemental Figure 4**). This raises the possibility that intracellular factors may be necessary for FAK to regulate PDE8A-driven cAMP hydrolysis in cells. Taken together, our findings that PDE8A inhibitors phenocopy the effects of FAK inhibitors in cells and that FAK can phosphorylate and associate with PDE8A suggest a model (**Figure 8**) in which PDE8A is a critical FAK substrate that participates in crosstalk between FAK and cAMP signaling.

**Figure 7.**
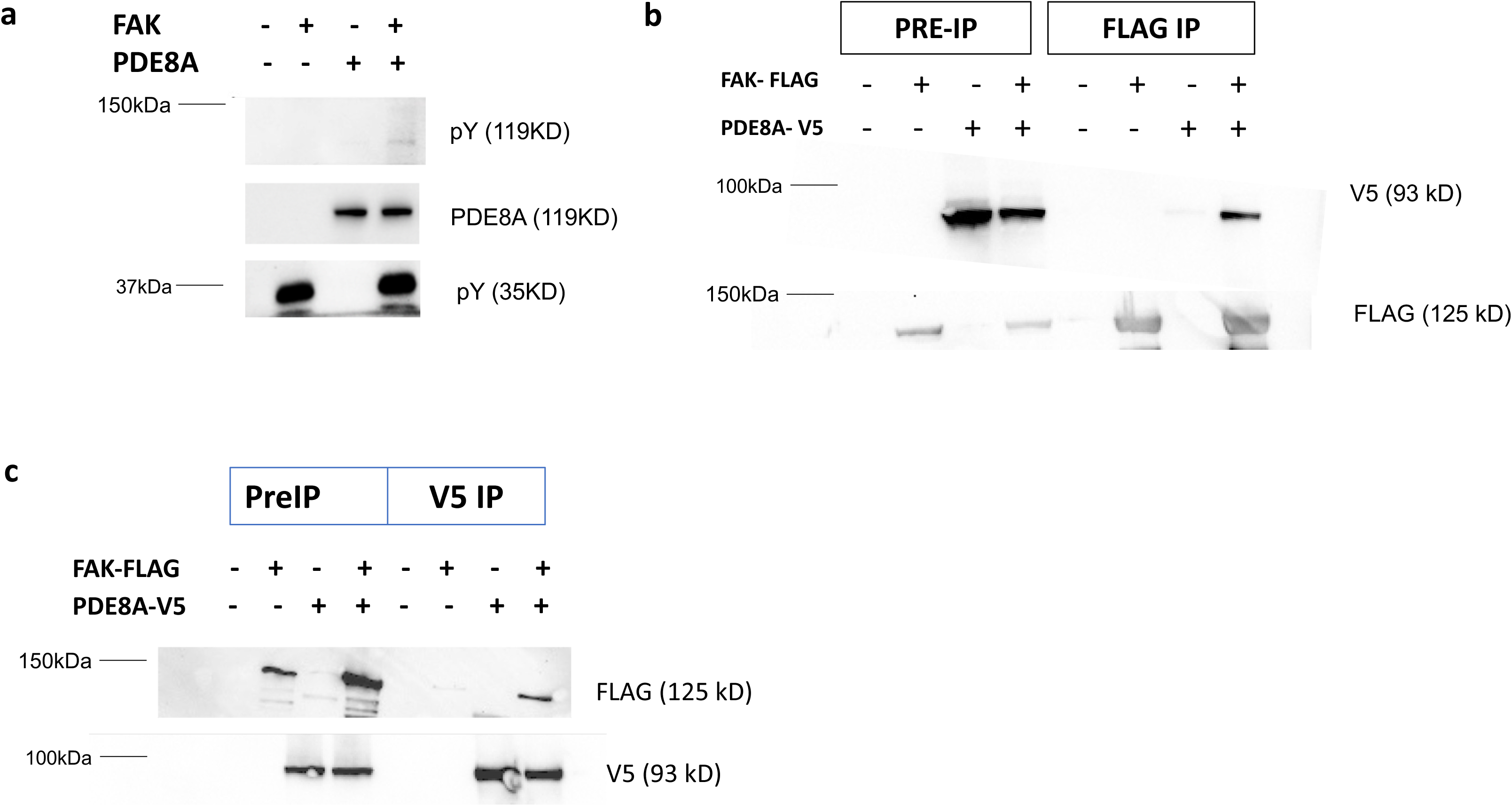
FAK phosphorylates PDE8A and there is an association between FAK and PDE8A in cells: (**a**) Immunoblotting for phospho-tyrosine (pY) and PDE8A *in vitro* FAK kinase assay with recombinant human PDE8A. pY bands are located at the expected size of FAK (auto-phosphorylation) and PDE8A. (**b**) Immunoblotting for FLAG and V5 in lysates from cells transfected with plasmids expressing FAK and PDE8A tagged with FLAG and V5 respectively after immunoprecipitation with anti-FLAG affinity gel. (**c**) Immunoblotting for FLAG and V5 in protein lysates from cells transfected with plasmids expressing FAK and PDE8A tagged with FLAG and V5 respectively after immunoprecipitation with anti-V5 affinity gel.

**Figure 8:**
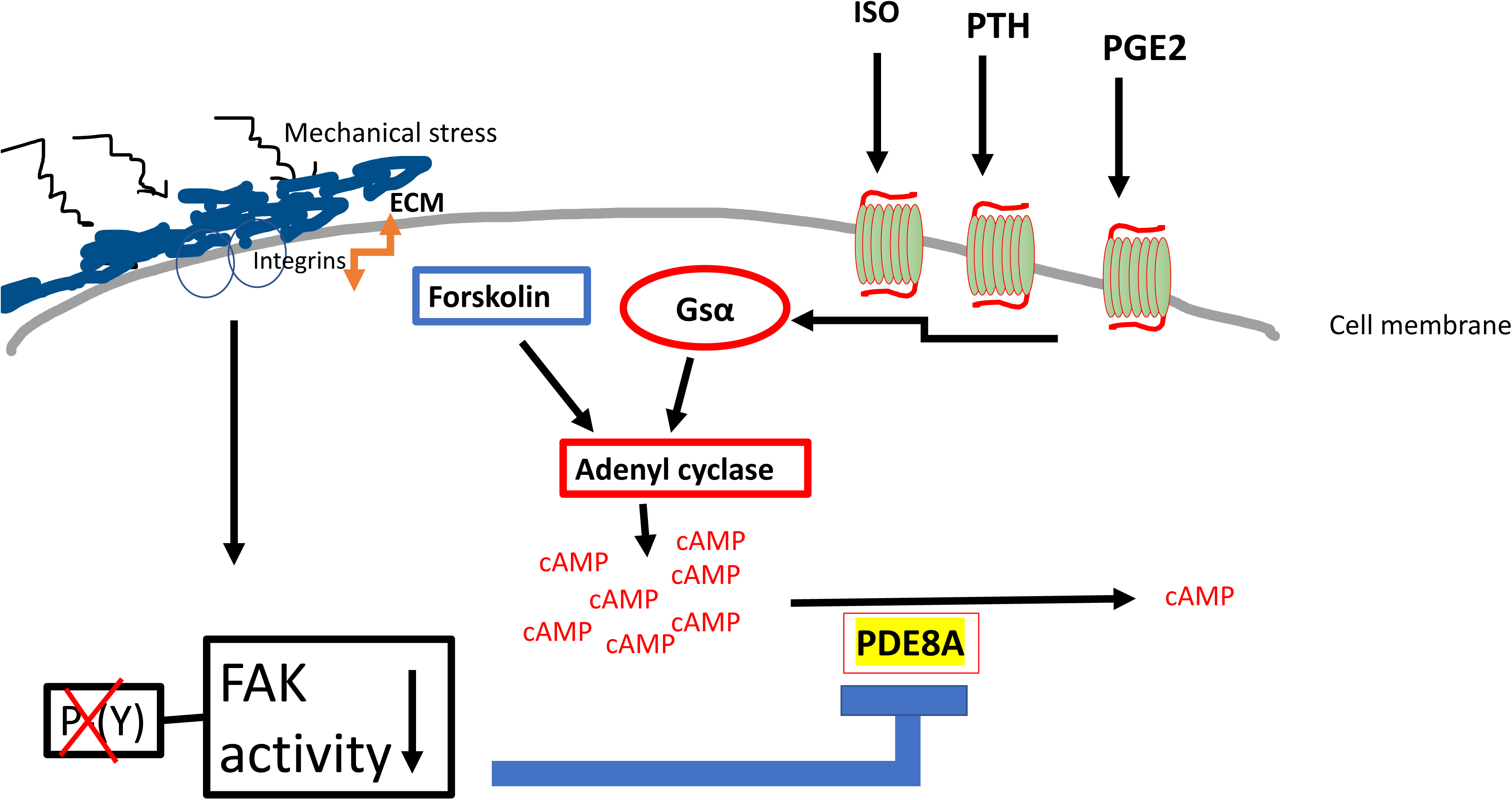
Model schematizing how FAK controls cAMP levels via PDE8A downstream of mechanical and hormonal cues. Normally, active FAK phosphorylates PDE8A and maintains high basal PDE8A-mediated cAMP hydrolysis. FFSS causes a reduction of FAK activity leading to reduced FAK-mediated PDE8A phosphorylation and, in turn, reduced PDE8A-mediated cAMP breakdown. Thus, FAK inhibition leads to suppressed PDE8A function causing an accumulation of cAMP that is generated by upstream GPCR signaling.

## Discussion

The present study suggests that the mechanosensitive FAK/PDE8A pathway enhances the effects of PTH signaling in osteocytes. FAK has an important role in cellular mechanotransduction, the process in which cells sense extracellular mechanical stimuli and translate them into chemical signals [34]. We previously demonstrated that osteocytes’ high basal FAK activity is inhibited by mechanical cues leading to reduction of its function. This subsequently leads to reduced tyrosine phosphorylation of class IIa HDACs, their nuclear translocation and suppression of MEF2-driven sclerostin expression [13]. Here, we find another link between mechanosensitive FAK and osteocyte function. Normally, active FAK phosphorylates PDE8A and maintains high basal PDE8A-mediated cAMP hydrolysis. Upon mechanical cues, FAK activity is temporarily suppressed, leading to reduced FAK-mediated PDE8A phosphorylation and, in turn, reduced PDE8A-mediated cAMP catabolism. Thus, FAK inhibition, either via mechanical signaling or by the direct FAK inhibition strategies used here, leads to suppressed PDE8A function causing an accumulation of cAMP (**Figure 8**). This is the first study to identify a direct link between mechanical stress, via FAK, and phosphodiesterase function. In addition, it is the first study to identify a role of PDE8A in osteocytes.

cAMP has a key role downstream of PTH to regulate bone remodeling [18–20]. PKA activation in human mesenchymal stromal cells increases osteogenesis [35]. It has also been suggested that the duration of cAMP signal is important in promoting or inhibiting osteogenesis [36] with transient cAMP signaling leading to osteogenesis whereas continuous cAMP blocks osteoblast maturation [37]. PTH generates cAMP through GPCR signaling in plasma membrane or within endosomes by PTH/PTH1R [38]. The cAMP signal strength and duration depend on the PTH concentration but also on the PTH1R conformation and the internalization of the receptor/ligand complex [18, 39]. Here, we report the regulation of cAMP levels by FAK-dependent PDE8A modulation as an additional signaling checkpoint after its generation in osteocytes.

The role of phosphodiesterases and the concept of using phosphodiesterase inhibitors to treat osteoporosis is not entirely new [37]. The inhibition of phosphodiesterases PDE2, PDE3 and PDE4 were found to increase osteoblast differentiation *in vitro* [40]. The administration of the non-specific PDE inhibitor pentoxifylline increased bone mass in mice through production of Wnt10b from CD8+ T cells [41]. Pentoxifylline also increased skeletal angiogenesis and reversed osteopenia in ovariectomized rats [42]. Pentoxifylline, when combined with PTH, enhanced the calcium content of BMP2- induced ectopic bone and an additive effect of PTH and phosphodiesterases in bone formation was suggested [43]. Both cAMP and cGMP phosphodiesterases have been shown to play a role in bone biology. The inhibition of the cGMP phosphodiesterase PDE5 increased angiogenesis and bone formation in a murine fracture model [44] and restored bone mass in ovariectomized mice [45]. The cAMP phosphodiesterase PDE4 inhibitor rolipram increased bone mass after 5 weeks of daily injections [46]. An association between bone mass and PDE4D gene polymorphisms was found in an English population sample [47]. Rolipram was found to have an inhibitory effect on osteoclastogenesis *in vivo* and *in vitro* [48, 49] though other *in vitro* studies suggested the opposite [50]. FAK associates with PDE4D in squamous cancer cells via an intermediate molecule, RACK1, to control cell polarization and adhesion [51]. This FAK/PDE4D interaction was also shown to play a role in melanoma cell invasion [52] and it is thought that via FAK/phosphodiesterase interaction the cell responds to signals from the extracellular environment [53]. Functional and physical associates between FAK and PDE8A have not been reported.

The predicted PDE8A tyrosine phosphorylation site that is regulated in response to FAK inhibition (Y315) is located in the PAS domain of the protein, a site of interaction with other proteins or molecules [54, 55]. Further study is needed to understand how FAK phosphorylation controls PDE8A activity in cells. Phosphodiesterase assays did not reveal obvious effects of FAK phosphorylation on PDE8A enzymatic activity *in vitro*. Since the predicted FAK phosphorylation site in PDE8A is outside of its phosphodiesterase domain, it is possible that FAK phosphorylation may promote PDE8A activity in cells via an allosteric mechanism requiring cofactors not present in our *in vitro* assays using recombinant proteins.

Future research will investigate the role of this novel pathway in bone. The effects of conditional FAK deletion *in vivo* appear to be cell stage-specific in the osteoblast lineage. When FAK was deleted in mice in osteoblast progenitors using Osx-Cre, decreased osteoblast number and low bone mass were observed [56]. A similar phenotype was seen by FAK deletion in osteoprogenitor cells using Dermo1-Cre [57]. Constitutive FAK deletion in mature osteoblasts via Col1a1-Cre yielded no obvious skeletal phenotype at baseline or in response to mechanical loading [58, 59]; however, adequate deletion of the FAK protein in bone cells was not shown. In addition, these mouse models demonstrate the results of continuous FAK inhibition in the bone cells. In contrast, daily injections of a dual FAK-PYK2 inhibitor caused net bone gain in an osteoporosis rat model [16]. PTH causes net bone gain when given intermittently via daily injections, while it causes net bone loss when it increases continuously such as in hyperparathyroidism [7]. FAK inhibition that, similar to PTH upregulates cAMP, may have discordant effects when it occurs continuously versus intermittently.

In summary, here we demonstrate a novel link between mechanical signals and phosphodiesterase function that regulates the cAMP pathway in osteocytes and results in enhanced parathyroid hormone signaling. These results provide a mechanistic connection between mechanical and PTH signaling in bone cells and suggest potential avenues for future development of osteoporosis therapeutics that simultaneously engage multiple bone anabolic pathways.

## Methods

### Phosphoproteomic analysis

Cell extracts from wild type Ocy454 cells, FAK KO Ocy454 cells and Ocy454 cells treated with FAK inhibitor (VS6063, at 10µM for 30 minutes) were prepared in Urea Lysis Buffer (9M Urea, 20mM HEPES pH 8.0 + phosphatase inhibitor cocktail) and shipped to Cell Signaling Technology for analysis. Cells were sonicated and centrifuged to remove insoluble material. Protein content was determined by Bradford assay and equal protein quantities from all samples were used for analysis. Samples were reduced with DTT and alkylated with iodoacetamide. Samples were digested with trypsin (CST #56296), purified over C18 columns (Waters #WAT051910) and enriched using the PTMScan Phosphotyrosine pY-1000 Kit (CST #8803) as previously described [60]. Samples were desalted over C18 tips prior to LC-MS/MS analysis. LC-MS/MS analysis was performed using a Thermo Orbitrap Fusion™ Lumos™ Tribrid™ mass spectrometer as previously described [60, 61] with replicate injections of each sample. Briefly, peptides were separated using a 50cm x 100µM PicoFrit capillary column packed with C18 reversed-phase resin and eluted with a 90-minute linear gradient of acetonitrile in 0.125% formic acid delivered at 280 nl/min. Tandem mass spectra were collected in a data-dependent manner using a 3 sec cycle time MS/MS method, a dynamic repeat count of one, and a repeat duration of 30 sec. Real time recalibration of mass error was performed using lock mass [62] with a singly charged polysiloxane ion m/z = 371.101237.

MS spectra were evaluated by Cell Signaling Technology using Comet and the GFY-Core platform (Harvard University) [63–65]. Searches were performed against the most recent update of the Uniprot *Mus musculus* database with a mass accuracy of +/-20 ppm for precursor ions and 0.02 Da for product ions. Cysteine carbamidomethylation was specified as a static modification, oxidation of methionine and serine, threonine, or tyrosine phosphorylation were allowed as variable modifications. Results were filtered to a 1% peptide-level FDR with mass accuracy +/-5ppm on precursor ions and presence of a phosphorylated residue for enriched samples. Site localization confidence was determined using AScore [66]. All quantitative results were generated using Skyline [67] to extract the integrated peak area of the corresponding peptide assignments. Accuracy of quantitative data was ensured by manual review in Skyline or in the ion chromatogram files. The relative abundance of each phosphorylated tyrosine residue was plotted in GraphPad prism. These results are shown in **Supplemental table 1.**

### Chemicals

Chemical reagents VS6063 (Selleckchem, S7654), PF431396 (Selleckchem, S7644), PF562271 (Selleckchem, S2890), GSK2256098 (MedChemExpress HY-100498), Y15 (MedChemExpress HY-12444), cilengitide (APExBIO A8660), PF04957325 (MedChemExpress HY-15426) were used at doses and times indicated in figure legends. All compounds were dissolved in DMSO as 10 mM stocks and stored at −20°C. Forskolin (Sigma F6886) was diluted in water with 2% ethanol as per manufacturer’s instructions. PTH (1-34) was synthesized by the Massachusetts General Hospital (MGH) peptide core facility.

### Cells and GloSensor assay

GS22 cells, HEK293 cells stably expressing the GloSensor cAMP reporter, GP2.3 cells, HEK293/GloSensor (GS22) cells stably expressing human PTH1R [24]; GBR-24 cells, HEK293/GloSensor (GS22) cells stably expressing βArrestin2^yfp^ [68][65]; GPG10 cells, HEK293/Glosensor (GS22) cells stably expressing a pH-sensitive PTH1R (PTH1R-GFP^pHs^) [68, 69], and GSG-5 cells, HEK293-Gαs-KO/GloSensor cells [70], were all maintained in DMEM (10% FBS, 1% penicillin/streptomycin) [71]. SGS-72 cells, human osteosarcoma SaOS-2 cells stably expressing the GloSensor cAMP reporter, were maintained in modified McCoy’s 5A medium (15% FBS, 1% penicillin/ streptomycin, and 1% nonessential amino acids) [23]. Cells were seeded into 96-well white plates (40,000 cells/100μL/well); 24 to 48 h after confluency, wells were rinsed with 100 μL carbon dioxide-independent culture medium (CIDB; Life Technologies) containing 0.1% bovine serum albumin (BSA), which was then replaced with 90 μL of luciferin (0.5 mM in CIDB) with FAK inhibitor or vehicle for 30 min before adding 10 μL of GPCR ligands (PTH 1-34, PGE_2_, Isoproterenol) or forskolin diluted in 10 μL of CIDB. cAMP-dependent luminescence was then measured at 2-min intervals using a PerkinElmer Envision 2104 plate reader (PerkinElmer), as described [24]. Ligand dose–response curves were generated as described [24]. Ocy454 cells were as previously described [26]. A single-cell sub- clone of Ocy454 cells was used for all experiments. Cells were seeded at 50,000 cells/mL and allowed to reach confluency at 33°C in 2–3 days. Then, cells were transferred from 33°C to 37°C to inactivate the temperature-sensitive T antigen and facilitate osteocytic differentiation. For protein and gene expression analyses, cells were analyzed after culture at 37°C for 14–21 days. Cells were routinely assessed for *Sost* and *Dmp1* expression at 37°C and examined for osteocytic morphology.

### CRISPR/Cas9-mediated gene deletion and lentiviral knockdown

To design sgRNA target sequences, we used “Design sgRNA for CRISPRko” web tool (https://portals.broadinstitute.org/gpp/public/analysis-tools/sgrna-design) and selected the top three sgRNA sequences. Target sequences were cloned into the PX458 vector which was subsequently transfected into GloSensor HEK293 cells using PolyJet transfection reagent (SignaGen SL100688). GFP high cells were recovered by flow cytometry based sorting at a density of one cell/well into 96 well plates. Single cell clones were selected and screened for FAK deletion by immunoblotting. FAK-KO clones 1–3 were used for subsequent experiments.

For shRNA experiments, HEK293 cells were transduced with lentiviruses harboring human PTK2 shRNAs that were cloned into the pLKO.1 backbone. To design the shRNA sequences we used https://portals.broadinstitute.org/gpp/public/gene/search and selected the top three sequences (clone IDs TRCN0000121318, TRCN0000196310, TRCN0000196667 – DNA sequences shown in Supplemental Table 3). The 3^rd^ sequence did not suppress FAK expression and was not used for subsequent experiments. To produce lentiviruses, HEK293T cells transfected with pLKO.1, psPAX2 (Addgene; plasmid #12260), and MD2.G (Addgene; plasmid #12259) using PolyJet transfection reagent (SignaGen SL100688). Medium was changed the next day, and then collected 48 h later. Cells were exposed to lentiviral particles overnight in the presence of polybrene (2.5 μg/mL). Media was then changed with puromycin (4 μg/mL). Cells were maintained in selection medium throughout the duration of the experiment (1 week). Control cells were transduced with an empty shRNA- expressing lentivirus.

### Immunoblotting

Whole-cell lysates for phosphorylated PKA substrate assessment were prepared using RIPA buffer (Boston BioProducts BP-115). Whole-cell lysates for immunoprecipitation assays were prepared using TNT (Tris-NaCl-Tween buffer, 20 mM Tris-HCl pH 8, 250 mM NaCl, 0.5% Triton X-100 containing protease inhibitor (PI), 1 mM NaF, 1 mM DTT, 1 mM vanadate). Adherent cells were washed with ice cold PBS, then scraped into RIPA or TNT buffer on ice. Material was then transferred into Eppendorf tubes kept on ice, vortexed at top speed for 30 s, then centrifuged at 14,000 g for 6 min at 4°C. The supernatants were collected, 2x SDS sample buffer was added and heated at 95°C for 5 min for immunoblotting. Lysates or immunoprecipitates were separated by SDS-PAGE, and proteins were transferred to the nitrocellulose (Amersam #10600008). Membranes were blocked with 5% milk in tris-buffered saline plus 0.05% Tween- 20 (TBST) and incubated with primary antibody overnight at 4°C. The next day, membranes were washed, incubated with appropriate HRP-coupled secondary antibodies, and signals detected with ECL Western Blotting Substrate (Pierce), ECL Plus Western Blotting Substrate (Pierce), or SuperSignal West Femto Maximum Sensitivity Substrate (Thermo scientific). Antibodies purchased from Cell Signalling Technology: phospho-PKA substrate (RRxS*/T*) (1:1000, Cell Signaling Technology, 9624S), FAK (1:1000, Cell Signaling Technology, 13009), p- FAK(Y397) (1:500, Cell Signaling Technology, 8556), beta-tubulin (1:250, Cell Signaling Technology, 5346), FLAG (1:1000, Cell Signaling Technology, 2368S), V5 (1:1000, Cell Signaling Technology, 13202S), phospho-tyrosine (1:1000, Cell Signaling Technology, 8954S)

### RNA isolation and RT-qPCR

Total RNA was collected from cultured cells using QIAshredder (QIAGEN) and PureLink RNA mini kit (Invitrogen) following the manufacturer’s instructions. In brief, lysis buffer with 2- mercaptoethanol was added to cold PBS-washed cells and collected into QIAshredder, then centrifuged at 15,000 g for 3 min. The flow-through was collected into a new tube, and RNA isolation was carried out with PureLink RNA mini kit following the manufacturer’s instructions. For qRT-PCR, cDNA was prepared with 750 ng of RNA using the Primescript RT kit (Takara Inc.) and analyzed with PerfeCa^®^ SYBR^®^ Green FAstMix^®^ ROX (Quanta bio) in the StepOnePlus^TM^ Real-time PCR System (Applied Biosystems) using specific primers designed for each targeted gene (**Supplemental Table 3**). Relative expression was calculated using the 2^−ΔΔCT^ method [72] by normalizing with β-actin housekeeping gene expression, and presented as relative abundance to β-actin.

### Plasmids

pLV[Exp]-Puro-EF1A>{rGs-WT-FLAG}, pLV[Exp]-Puro-EF1A>{rGs-WT-FLAG-YFYF}, pRP- hP1R, pRP[Exp]-CMV>{hP1R-HA-Y494F,Y496F}, pRP-EF1a, pRP- EF1a> V5/hPDE8A and pRP EF1A> FLAG/hFAK were synthesized by VectorBuilder.

### PTH1R ß-arrestin recruitment, internalization, and intracellular calcium flux assays

Recruitment of β*-*arrestin was assessed in GBR-24 cells, HEK293/GloSensor (GS22) cells stably expressing βArrestin2^yfp^ [68, 69] and transiently transfected to express PTH1R-WT or PTH1R-Y494F,Y496F [73]. Cells were treated with red fluorescent PTH(1-34)^tmr^ and imaged by fluorescent microscopy.

Internalization was measured in GPG10 cells (HEK293/glosensor/PTH1R-GFPpH stable expression) as the change in the ratio of fluorescence derived from PTH1R-GFP^pHs^ upon sequential excitation at wavelengths (λ_ex_) of 485 and 405 nm and fluorescence emission at a wavelength (λ_em_) of 535 nm. Peptides were added to cells in black-walled 96-well plates and fluorescence was recorded at 1-minute plate intervals over the course of 90 minutes [73].

Signaling via the intracellular Ca^++^ pathway was assessed in stably transfected HEK- HEK293/hPTH1R/glosensor (GP-2.3) cells using the calcium-sensitive fluorophore Fura2-AM (Invitrogen, Life Tech. Grand Island, NY, USA) [73, 74]. Confluent cells in a black, 96-well plate were preloaded with Fura2-AM (5μM) for 45 min and then unloaded in buffer for 30 min. The plate was then processed using a Perkin Elmer Envision plate reader (Perkin Elmer, Waltham, MA, USA) to monitor fluorescence emission at a wavelength (λ_em_) of 515 nm, upon sequential excitation at wavelengths (λ_ex_) of 340 nm and 380 nm. Data were recorded in each single well at 2-s intervals for 10 s prior to, and for 150 s after ligand addition. The data at each time point were calculated as the ratio of the fluorescence signal obtained with excitation at 340 nm to that obtained with excitation at 380 nm.

### FAK kinase assay

Recombinant human PDE8A (Sigma SRP0273) was incubated with 100 ng FAK tyrosine kinase (Promega #V1971) 50μM DTT, 2mM MnCl_2_, 50μM ATP and 1x kinase buffer for 15 min at 37°C in a reaction of total volume 25 µL. The kinase reacted protein samples were boiled with 2- mercaptoethanol containing sample buffer for 5 min and loaded into 8% polyacrylamide gels for immunoblotting with phospho-tyrosine and PDE8A antibodies.

### Co-immunoprecipitation (co-IP) assays

For the co-IP experiments in 293T cells, a total of 5 μg of the following plasmids were transfected using PolyJet transfection reagent (SignaGen SL100688): EF1A empty vector, EF1A-FLAG/hFAK, EF1A-V5/hPDE8A plasmid. Forty-eight hours later, protein lysates were collected with 1 mL of TNT (Tris-NaCl-Tween buffer, 20 mM Tris-HCl pH 8, 200 mM NaCl, 0.5% Triton X-100 containing protease inhibitors (PI), 1 mM NaF, 1 mM DTT, 1 mM vanadate). 10% of the protein lysate was saved as a “pre-IP” sample. The remaining protein lysate was incubated with 25 μl of anti-FLAG affinity gel (Selleckchem B23101), rotated at 4°C overnight after which it was washed twice with TNT buffer containing PI, NaF, DTT, and vanadate. Precipitated samples were eluted using FLAG peptide 1μg/μl in elution buffer (50mM Tris PH 7.5, 150mM NaCl) supplemented with protease inhibitors (PI), 1 mM NaF, 1 mM DTT, 1 mM vanadate. After centrifugation at 5000 rpm for 1 min, the supernatants were collected for western blot analysis (IP samples). In both pre-IP and IP samples 50 μL SDS sample buffer was added and heated at 95°C for 5 min. In separate experiments, protein lysates were incubated with V5 affinity gel (Millipore A7345) and eluted with V5 peptide (Sigma Aldrich V7754) and with similar remaining steps.

### PDE8A Activity assay

PDE8A activity assay was performed with recombinant PDE8A and cAMP detection using the LANCE *Ultra* cAMP kit (Perkin Elmer). To determine the most sensitive range for the assay, the concentration of the cAMP standard supplied in the kit was titrated. The TR-FRET signal at 665 nm was normalized to the signal of the donor-channel at 620 nm to correct for well-to-well variability of the signal. A linear working cAMP range was obtained between 0.5 and 10 nM. Therefore, a cAMP concentration of 6 nM was used for subsequent assays. To optimize assay conditions, the effect of enzyme concentration on cAMP levels was first examined. Different concentrations (between 0.003 and 1 ng/µL) PDE8A (Sigma SRP0273) were tested in the presence of 6 nM cAMP in buffer containing 1X HBSS, 5 mM HEPES, 0.1% BSA, 3mM MgCl_2_, 50μM ATP. The reaction was stopped after 1 hour of incubation at RT by the addition of stop/detection buffer containing 1 mM IBMX and additional incubation for 1 hour at RT. The signal was read on the Envision plate reader 2104 at 620 and 665 nm. The assay was performed in white, opaque 384-well microplates. We noted submaximal but detectable PDE8A- stimulated cAMP hydrolysis when using PDE8A concentrations between 0.01-0.1 ng/µL for 1h. Consequently, 0.1 ng/µL PDE8A and an incubation time of one hour were selected for further experiments. Recombinant PDE8A and FAK (1 ng) proteins were added together in the reaction buffer and 665/620 ratios were measured after the above procedure.

### PDE profiling

PDE profiling of VS6063 against 13 PDEs was performed by Reaction Biology VS6063 was provided at a concentration of 10mM in DMSO. VS6063 was tested in a 10-dose IC50, singlet, with 3-fold serial dilutions starting at 10 μM. Control compounds were tested in a 10-dose IC50, singlet, with 3-fold serial dilution starting at different concentrations.

For PDE3A, PDE3B, PDE4A, PDE4B, PDE4C, PDE4D, PDE7A, PDE8A and PDE10A the reaction product, AMP and for PDE1A, PDE1B, PDE1C, and PDE2A the reaction product, GMP were detected by Transcreener fluorescent polarization assay.

Subtrate concentrations used: 1 μM cAMP for PDE3A, PDE3B, PDE4A, PDE4B, PDE4C, PDE4D, and PDE10A, 0.2 μM cAMP for PDE7A and PDE8A, 1 μM cGMP for PDE1A, PDE1B, PDE1C, and PDE2A. The reaction was performed at room temperature for 1hr. The assay buffer contained 10 mM Tris, pH 7.5, 5 mM MgCl2, 0.01% Brij 35, 1 mM DTT, and 1% DMSO and for PDE1A, PDE1B and PDE1C, 0.2 mM CaCl2 and 0.36 uM CaM were also added.

Compounds in DMSO were added into enzyme using Acoustic Technology. Substrate was added and incubated for 1 hour. The reaction was stopped with the addition of Stop/detection mixture (40 mM Tris-HCL, pH 7.5, containing 80 mM EDTA and 0.04% Brij35). Fluorescent polarization was measured after 90 minute incubation at room temperature and calculated mP (Ex = 620 nm FP, Em = 688 nm P and S). Data was analyzed based on AMP or GMP standard curve to obtain product, AMP or GMP, amount.

### Statistical analysis

All statistical analyses were performed by GraphPad Prism 10 for Mac (GraphPad Software Inc, USA). For comparison between two experimental groups, two-tailed t tests were used. When more than two groups were present, ANOVA with posthoc Tukey test was used. Values were expressed as mean ± SEM unless otherwise stated. A p-value < 0.05 was considered significant.

## Acknowledgements

GP acknowledges support from the MGH Endocrine Division T32 NIH training grant T32DK007028 and the Endocrine Fellows Foundation. MNW acknowledges support from DK116716, the Smith Family Foundation Odyssey Award, and a MGH Research Scholar award. TJG acknowledges support from DK11479, DK113039, and AR066261. TS acknowledges support from NIH/NIAMS (R21AR079633) and Center for Skeletal Research P30 core P&F award.

## Figure legends

**Supplemental Figure 1:**
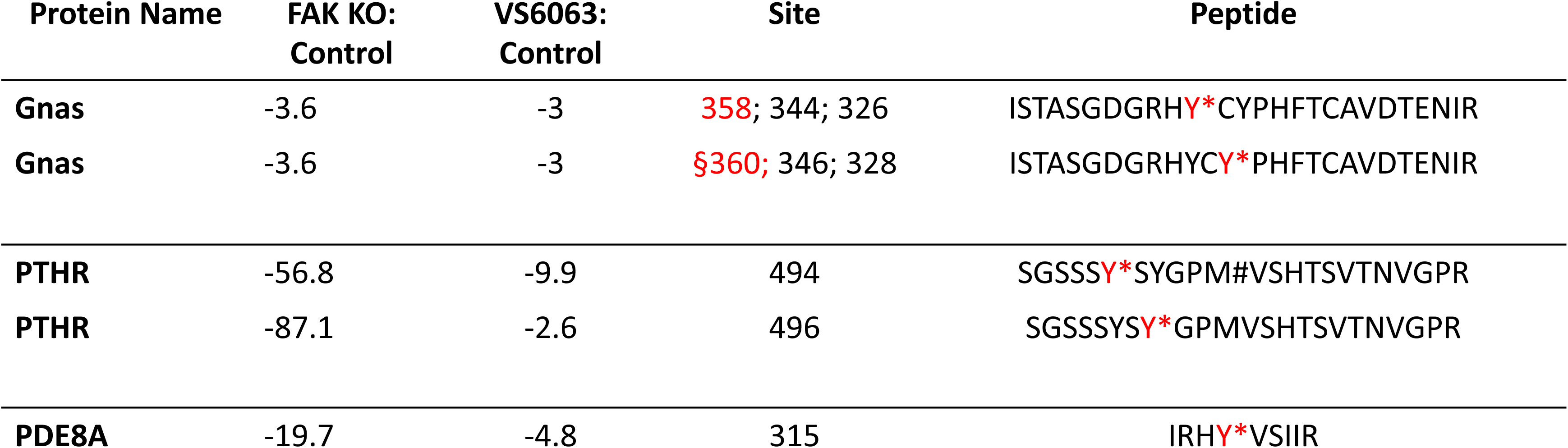
Gsα, PTHR1 and PDE8A tyrosine phosphorylation sites identified by phosphoproteomics to be reduced by FAK inhibition or deletion.

**Supplemental Figure 2:**
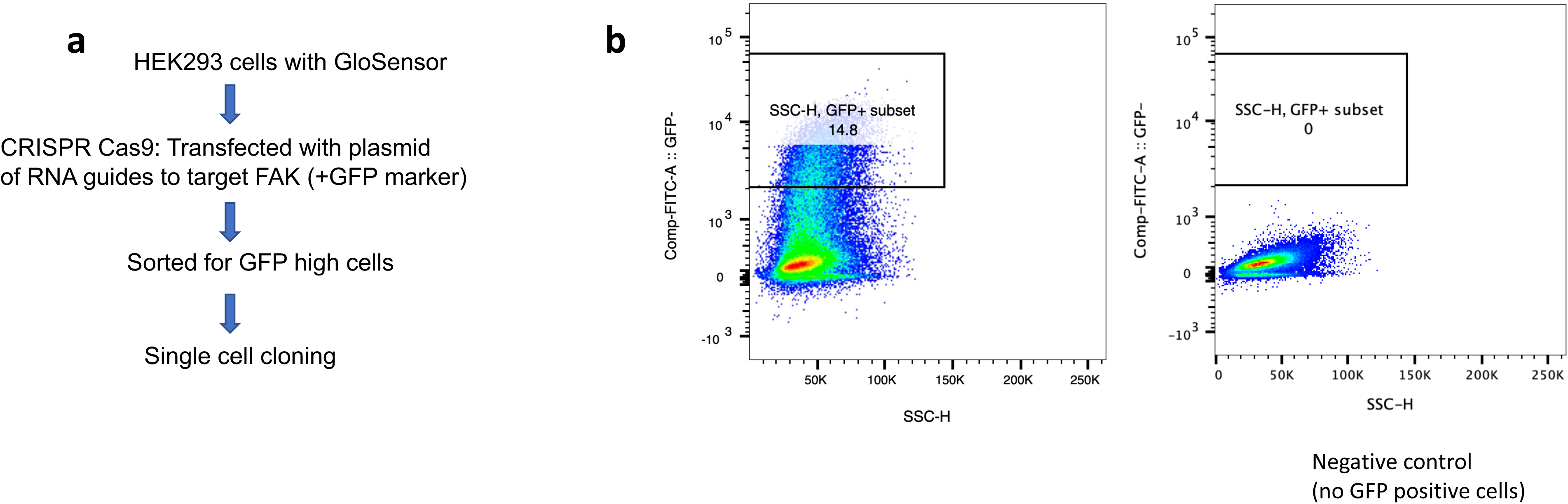
(**a**) **Strategy for deletion of FAK in GloSensor HEK cells** (**b**) Gating/sorting strategy. GFP high cells were single sorted to generate FAK KO single cell clones. Cells not transfected with plasmid harboring GFP marker are shown on the right.

**Supplemental Figure 3:**
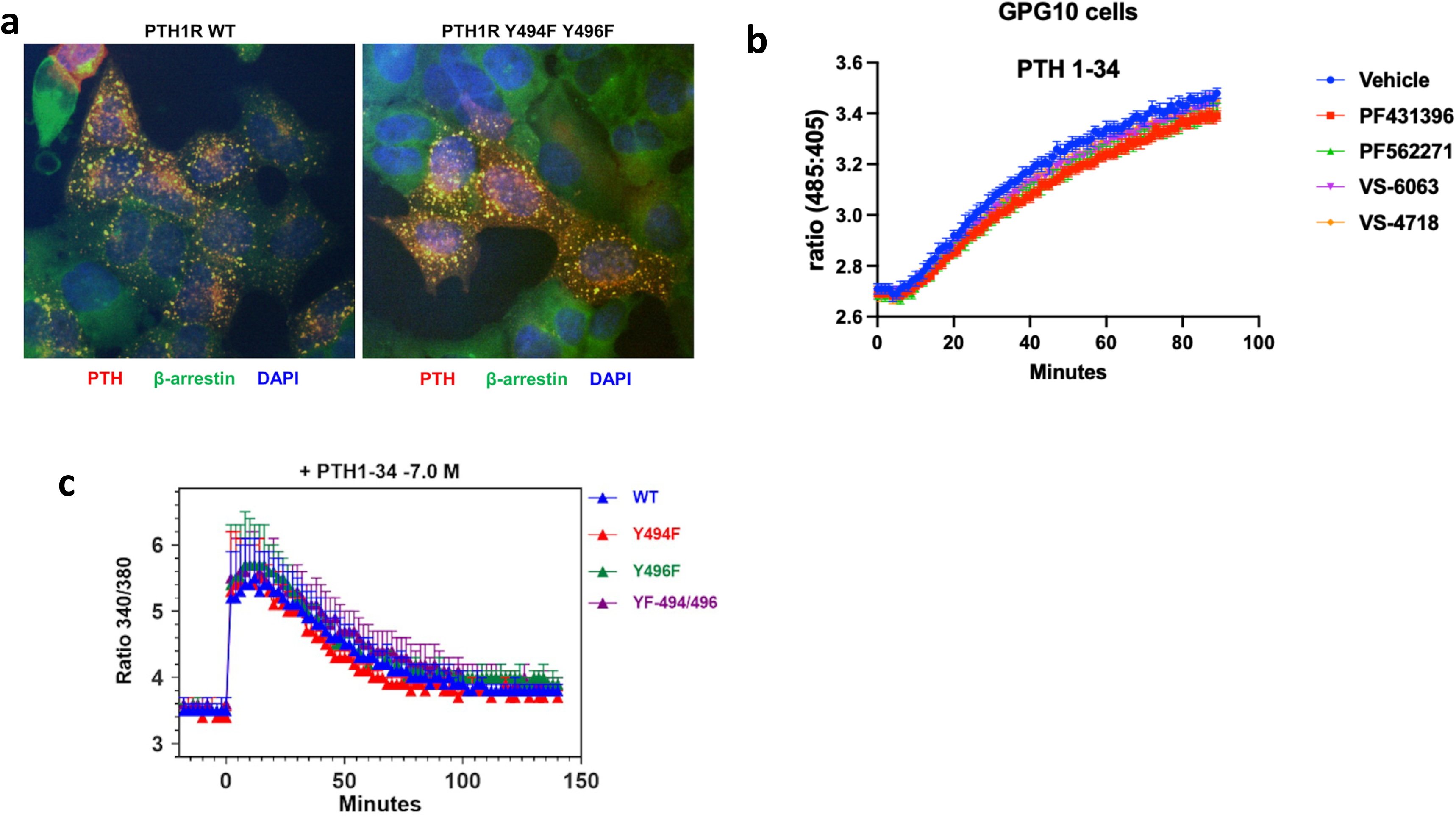
FAK inhibition does not influence PTH-stimulated PTH1R ß- arrestin recruitment, internalization, or intracellular calcium signaling. **(a)** The βarrestin2yfp clustering and colocalizing with PTH(1-34)^tmr^ was induced similarly between the WT PTH1R and the mutant Y494F and Y496F PTH1R in GBR-24 cells. **(b)** HEK 293 cells stably expressing PTH1R-GFP^pHs^ (GPG-10 cells) were treated with PTH 1-34 and 10μM of the FAK inhibitors for 90 min. Internalization was measured as an increase in the ratio of GFP^pHs^ fluorescence at a wavelength (λ_em_) of 535 nm upon sequential excitation at wavelengths (λ_ex_) of 485 and 405 nm; data are means±SEM from n=6 independent experiments. **(c)** HEK 293 cells (hPTH1R/GloSensor (GP-2.3 cells) were preloaded with the calcium indicator dye Fura2-AM, and then treated with PTH 1-34 (100 nM). Intracellular calcium fluxes were measured by ratiometric fluorescence (excitation = 340 nm and 380 nm, emission = 535 nm) over time.

**Supplemental Figure 4:**
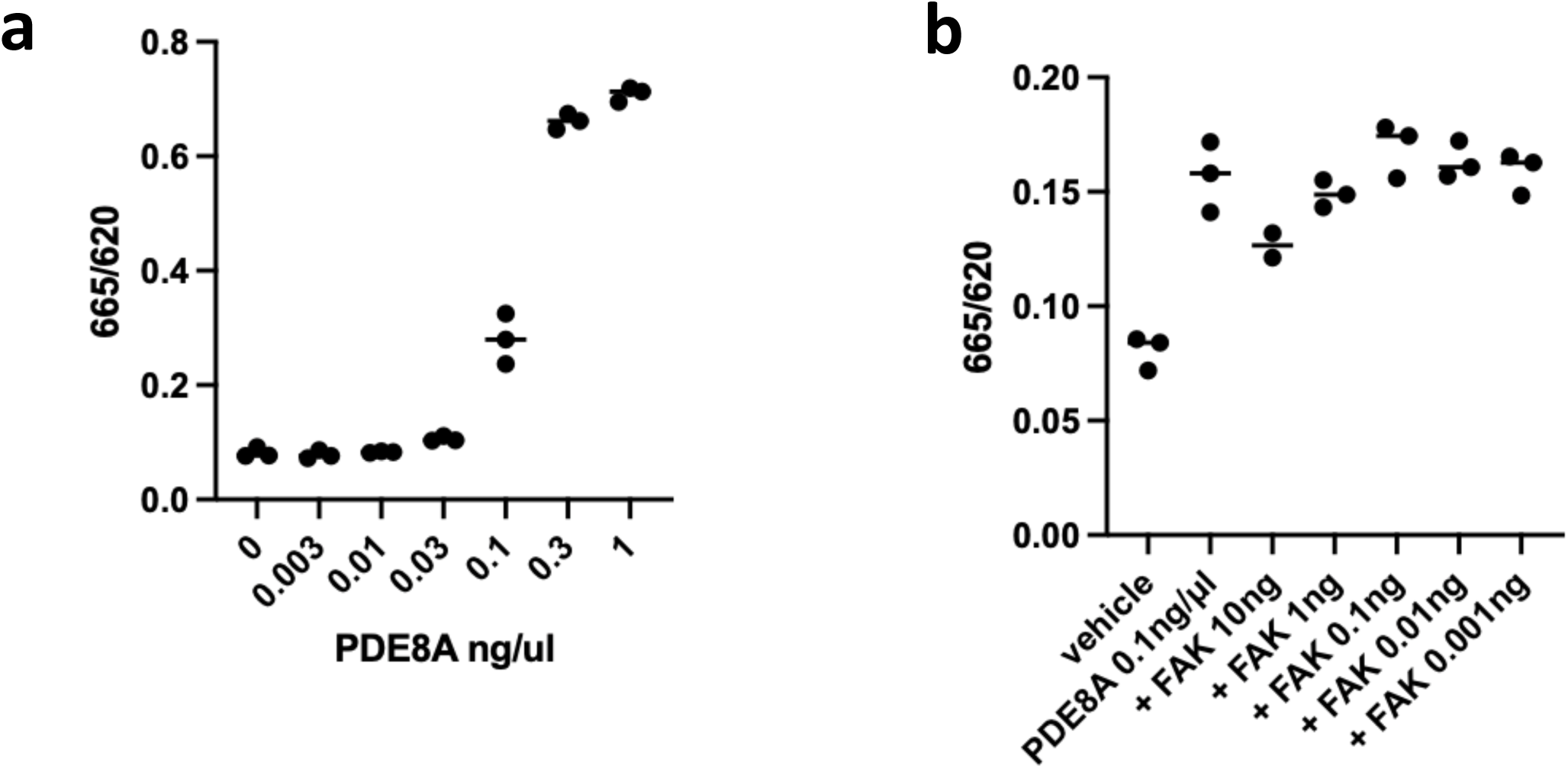
PDE8A activity assay: FAK does not increase the activity of PDE8A in a cell free system. (**a**) PDE8A dose response. X axis: recombinant PDE8A dose in ng/μl y axis: 665/620 TR-FRET signal ratio (inversely proportional to cAMP levels) (**b**) 665/620 ratio after incubation of recombinant PDE8A with different amounts of recombinant FAK (shown on axis x). All reactions occurred in presence of 6 nM cAMP. TR-FRET signal at 665 nm was normalized to the signal of the donor-channel at 620 nm to correct for well-to-well variability of the signal

**Supplemental Table 1:** Full data report from Phosphoproteomic assay

**Supplemental Table 2:** VS6063 tested *in vitro* against a panel of recombinant phosphodiesterases. Up to 10 µM VS6063 concentrations did not significantly inhibit PDE activity except a weak effect on PDE2A.

**Supplemental Table 3:** List of nucleotide sequences of primers, CRISPR guides and shRNA sequences.

## References

1. Hamilton, C.J., V.J. Swan, and S.A. Jamal, The effects of exercise and physical activity participation on bone mass and geometry in postmenopausal women: a systematic review of pQCT studies. Osteoporos Int, 2010. 21(1): p. 11–23.

2. Antoniou, G., et al., Bone Mineral Density Post a Spinal Cord Injury: A Review of the Current Literature Guidelines. Cureus, 2022. 14(3): p. e23434.

3. Wein, M.N., Parathyroid Hormone Signaling in Osteocytes. JBMR Plus, 2018. 2(1): p. 22–30.

4. Li, M.C.M., et al., The role of osteocytes-specific molecular mechanism in regulation of mechanotransduction - A systematic review. J Orthop Translat, 2021. 29: p. 1–9.

5. Palumbo, C. and M. Ferretti, The Osteocyte: From “Prisoner” to “Orchestrator”. J Funct Morphol Kinesiol, 2021. 6(1).

6. Robling, A.G., et al., Mechanical stimulation of bone in vivo reduces osteocyte expression of Sost/sclerostin. J Biol Chem, 2008. 283(9): p. 5866–75.

7. Silva, B.C. and J.P. Bilezikian, Parathyroid hormone: anabolic and catabolic actions on the skeleton. Curr Opin Pharmacol, 2015. 22: p. 41–50.

8. Turner, R.T., et al., Disuse in adult male rats attenuates the bone anabolic response to a therapeutic dose of parathyroid hormone. J Appl Physiol (1985), 2006. 101(3): p. 881–6.

9. Tanaka, S., et al., Skeletal unloading alleviates the anabolic action of intermittent PTH(1- 34) in mouse tibia in association with inhibition of PTH-induced increase in c-fos mRNA in bone marrow cells. J Bone Miner Res, 2004. 19(11): p. 1813–20.

10. Kostenuik, P.J., et al., Skeletal unloading causes resistance of osteoprogenitor cells to parathyroid hormone and to insulin-like growth factor-I. J Bone Miner Res, 1999. 14(1): p. 21–31.

11. Gardinier, J.D., F. Mohamed, and D.H. Kohn, PTH Signaling During Exercise Contributes to Bone Adaptation. J Bone Miner Res, 2015. 30(6): p. 1053–63.

12. Jepsen, D.B., et al., The combined effect of Parathyroid hormone (1-34) and whole-body Vibration exercise in the treatment of postmenopausal OSteoporosis (PaVOS study): a randomized controlled trial. Osteoporos Int, 2019. 30(9): p. 1827–1836.

13. Sato, T., et al., A FAK/HDAC5 signaling axis controls osteocyte mechanotransduction. Nat Commun, 2020. 11(1): p. 3282.

14. Geiger, B., J.P. Spatz, and A.D. Bershadsky, Environmental sensing through focal adhesions. Nat Rev Mol Cell Biol, 2009. 10(1): p. 21–33.

15. Tilghman, R.W. and J.T. Parsons, Focal adhesion kinase as a regulator of cell tension in the progression of cancer. Semin Cancer Biol, 2008. 18(1): p. 45–52.

16. Buckbinder, L., et al., Proline-rich tyrosine kinase 2 regulates osteoprogenitor cells and bone formation, and offers an anabolic treatment approach for osteoporosis. Proc Natl Acad Sci U S A, 2007. 104(25): p. 10619–24.

17. Kang, Y., et al., Role of focal adhesion kinase in regulating YB-1-mediated paclitaxel resistance in ovarian cancer. J Natl Cancer Inst, 2013. 105(19): p. 1485–95.

18. Sutkeviciute, I., et al., PTH/PTHrP Receptor Signaling, Allostery, and Structures. Trends Endocrinol Metab, 2019. 30(11): p. 860–874.

19. Sinha, P., et al., Loss of Gsalpha early in the osteoblast lineage favors adipogenic differentiation of mesenchymal progenitors and committed osteoblast precursors. J Bone Miner Res, 2014. 29(11): p. 2414–26.

20. Wu, J.Y., et al., Gsalpha enhances commitment of mesenchymal progenitors to the osteoblast lineage but restrains osteoblast differentiation in mice. J Clin Invest, 2011. 121(9): p. 3492–504.

21. Buccioni, M., et al., Innovative functional cAMP assay for studying G protein-coupled receptors: application to the pharmacological characterization of GPR17. Purinergic Signal, 2011. 7(4): p. 463–8.

22. Hanna, P., et al., Homozygous Ser-1 to Pro-1 mutation in parathyroid hormone identified in hypocalcemic patients results in secretion of a biologically inactive pro-hormone. Proc Natl Acad Sci U S A, 2023. 120(8): p. e2208047120.

23. Lee, S., et al., A Homozygous [Cys25]PTH(1-84) Mutation That Impairs PTH/PTHrP Receptor Activation Defines a Novel Form of Hypoparathyroidism. J Bone Miner Res, 2015. 30(10): p. 1803–13.

24. Guo, J., et al., Prolonged Pharmacokinetic and Pharmacodynamic Actions of a Pegylated Parathyroid Hormone (1-34) Peptide Fragment. J Bone Miner Res, 2017. 32(1): p. 86–98.

25. Kapp, T.G., et al., A Comprehensive Evaluation of the Activity and Selectivity Profile of Ligands for RGD-binding Integrins. Sci Rep, 2017. 7: p. 39805.

26. Spatz, J.M., et al., The Wnt Inhibitor Sclerostin Is Up-regulated by Mechanical Unloading in Osteocytes in Vitro. J Biol Chem, 2015. 290(27): p. 16744–58.

27. Zhang, P., et al., Structure and allostery of the PKA RIIbeta tetrameric holoenzyme. Science, 2012. 335(6069): p. 712–6.

28. Zhao, L.H., et al., Structure and dynamics of the active human parathyroid hormone receptor-1. Science, 2019. 364(6436): p. 148–153.

29. Qiu, T., et al., IGF-I induced phosphorylation of PTH receptor enhances osteoblast to osteocyte transition. Bone Res, 2018. 6: p. 5.

30. Wein, M.N. and H.M. Kronenberg, Regulation of Bone Remodeling by Parathyroid Hormone. Cold Spring Harb Perspect Med, 2018. 8(8).

31. Cheloha, R.W., et al., PTH receptor-1 signalling-mechanistic insights and therapeutic prospects. Nat Rev Endocrinol, 2015. 11(12): p. 712–24.

32. Fisher, D.A., et al., Isolation and characterization of PDE8A, a novel human cAMP-specific phosphodiesterase. Biochem Biophys Res Commun, 1998. 246(3): p. 570–7.

33. Basole, C.P., et al., Treatment of Experimental Autoimmune Encephalomyelitis with an Inhibitor of Phosphodiesterase-8 (PDE8). Cells, 2022. 11(4).

34. Urciuoli, E. and B. Peruzzi, Involvement of the FAK Network in Pathologies Related to Altered Mechanotransduction. Int J Mol Sci, 2020. 21(24).

35. Siddappa, R., et al., cAMP/PKA pathway activation in human mesenchymal stem cells in vitro results in robust bone formation in vivo. Proc Natl Acad Sci U S A, 2008. 105(20): p. 7281–6.

36. Siddappa, R., et al., Timing, rather than the concentration of cyclic AMP, correlates to osteogenic differentiation of human mesenchymal stem cells. J Tissue Eng Regen Med, 2010. 4(5): p. 356–65.

37. Porwal, K., et al., Therapeutic potential of phosphodiesterase inhibitors in the treatment of osteoporosis: Scopes for therapeutic repurposing and discovery of new oral osteoanabolic drugs. Eur J Pharmacol, 2021. 899: p. 174015.

38. Vilardaga, J.P., F.G. Jean-Alphonse, and T.J. Gardella, Endosomal generation of cAMP in GPCR signaling. Nat Chem Biol, 2014. 10(9): p. 700–6.

39. Pena, K.A., S. Savransky, and B. Lewis, Endosomal signaling via cAMP in parathyroid hormone (PTH) type 1 receptor biology. Mol Cell Endocrinol, 2024. 581: p. 112107.

40. Wakabayashi, S., et al., Involvement of phosphodiesterase isozymes in osteoblastic differentiation. J Bone Miner Res, 2002. 17(2): p. 249–56.

41. Roser-Page, S., et al., Cyclic Adenosine Monophosphate (cAMP)-Dependent Phosphodiesterase Inhibition Promotes Bone Anabolism Through CD8(+) T Cell Wnt- 10b Production in Mice. JBMR Plus, 2022. 6(7): p. e10636.

42. Pal, S., et al., Reversal of Osteopenia in Ovariectomized Rats by Pentoxifylline: Evidence of Osteogenic and Osteo-Angiogenic Roles of the Drug. Calcif Tissue Int, 2019. 105(3): p. 294–307.

43. Horiuchi, H., et al., Enhancement of recombinant human bone morphogenetic protein-2 (rhBMP-2)-induced new bone formation by concurrent treatment with parathyroid hormone and a phosphodiesterase inhibitor, pentoxifylline. J Bone Miner Metab, 2004. 22(4): p. 329–34.

44. Menger, M.M., et al., Sildenafil, a phosphodiesterase-5 inhibitor, stimulates angiogenesis and bone regeneration in an atrophic non-union model in mice. J Transl Med, 2023. 21(1): p. 607.

45. Pal, S., et al., Skeletal restoration by phosphodiesterase 5 inhibitors in osteopenic mice: Evidence of osteoanabolic and osteoangiogenic effects of the drugs. Bone, 2020. 135: p. 115305.

46. Kinoshita, T., et al., Phosphodiesterase inhibitors, pentoxifylline and rolipram, increase bone mass mainly by promoting bone formation in normal mice. Bone, 2000. 27(6): p. 811–7.

47. Reneland, R.H., et al., Association between a variation in the phosphodiesterase 4D gene and bone mineral density. BMC Med Genet, 2005. 6: p. 9.

48. Miyamoto, K., et al., Phosphodiesterase 4 inhibitor rolipram potentiates the inhibitory effect of calcitonin on osteoclastogenesis. J Bone Miner Metab, 2006. 24(4): p. 260–5.

49. Park, H. and M. Yim, Rolipram, a phosphodiesterase 4 inhibitor, suppresses PGE2- induced osteoclast formation by lowering osteoclast progenitor cell viability. Arch Pharm Res, 2007. 30(4): p. 486–92.

50. Takami, M., et al., Phosphodiesterase inhibitors stimulate osteoclast formation via TRANCE/RANKL expression in osteoblasts: possible involvement of ERK and p38 MAPK pathways. FEBS Lett, 2005. 579(3): p. 832–8.

51. Serrels, B., et al., A complex between FAK, RACK1, and PDE4D5 controls spreading initiation and cancer cell polarity. Curr Biol, 2010. 20(12): p. 1086–92.

52. Delyon, J., et al., PDE4D promotes FAK-mediated cell invasion in BRAF-mutated melanoma. Oncogene, 2017. 36(23): p. 3252–3262.

53. Yarwood, S.J., E. Parnell, and R.J. Bird, The cyclic AMP phosphodiesterase 4D5 (PDE4D5)/receptor for activated C-kinase 1 (RACK1) signalling complex as a sensor of the extracellular nano-environment. Cell Signal, 2017. 35: p. 282–289.

54. Gu, Y.Z., J.B. Hogenesch, and C.A. Bradfield, The PAS superfamily: sensors of environmental and developmental signals. Annu Rev Pharmacol Toxicol, 2000. 40: p. 519–61.

55. Soderling, S.H., S.J. Bayuga, and J.A. Beavo, Cloning and characterization of a cAMP-specific cyclic nucleotide phosphodiesterase. Proc Natl Acad Sci U S A, 1998. 95(15): p. 8991–6.

56. Sun, C., et al., FAK Promotes Osteoblast Progenitor Cell Proliferation and Differentiation by Enhancing Wnt Signaling. J Bone Miner Res, 2016. 31(12): p. 2227–2238.

57. Qi, S., et al., FAK Promotes Early Osteoprogenitor Cell Proliferation by Enhancing mTORC1 Signaling. J Bone Miner Res, 2020. 35(9): p. 1798–1811.

58. Castillo, A.B., et al., Focal adhesion kinase plays a role in osteoblast mechanotransduction in vitro but does not affect load-induced bone formation in vivo. PLoS One, 2012. 7(9): p. e43291.

59. Kim, J.B., et al., Reconciling the roles of FAK in osteoblast differentiation, osteoclast remodeling, and bone regeneration. Bone, 2007. 41(1): p. 39–51.

60. Stokes, M.P., et al., Complementary PTM Profiling of Drug Response in Human Gastric Carcinoma by Immunoaffinity and IMAC Methods with Total Proteome Analysis. Proteomes, 2015. 3(3): p. 160–183.

61. Possemato, A.P., et al., Multiplexed Phosphoproteomic Profiling Using Titanium Dioxide and Immunoaffinity Enrichments Reveals Complementary Phosphorylation Events. J Proteome Res, 2017. 16(4): p. 1506–1514.

62. Olsen, J.V., et al., Parts per million mass accuracy on an Orbitrap mass spectrometer via lock mass injection into a C-trap. Mol Cell Proteomics, 2005. 4(12): p. 2010–21.

63. Eng, J.K., T.A. Jahan, and M.R. Hoopmann, Comet: an open-source MS/MS sequence database search tool. Proteomics, 2013. 13(1): p. 22–4.

64. Huttlin, E.L., et al., A tissue-specific atlas of mouse protein phosphorylation and expression. Cell, 2010. 143(7): p. 1174–89.

65. Villen, J., et al., Large-scale phosphorylation analysis of mouse liver. Proc Natl Acad Sci U S A, 2007. 104(5): p. 1488–93.

66. Beausoleil, S.A., et al., A probability-based approach for high-throughput protein phosphorylation analysis and site localization. Nat Biotechnol, 2006. 24(10): p. 1285–92.

67. MacLean, B., et al., Skyline: an open source document editor for creating and analyzing targeted proteomics experiments. Bioinformatics, 2010. 26(7): p. 966–8.

68. Cheloha, R.W., et al., Improved GPCR ligands from nanobody tethering. Nat Commun, 2020. 11(1): p. 2087.

69. Wehbi, V.L., et al., Noncanonical GPCR signaling arising from a PTH receptor-arrestin- Gbetagamma complex. Proc Natl Acad Sci U S A, 2013. 110(4): p. 1530–5.

70. Biebermann, H., et al., A New Multisystem Disorder Caused by the Galphas Mutation p.F376V. J Clin Endocrinol Metab, 2019. 104(4): p. 1079–1089.

71. Maeda, A., et al., Critical role of parathyroid hormone (PTH) receptor-1 phosphorylation in regulating acute responses to PTH. Proc Natl Acad Sci U S A, 2013. 110(15): p. 5864–9.

72. Livak, K.J. and T.D. Schmittgen, Analysis of relative gene expression data using real-time quantitative PCR and the 2(-Delta Delta C(T)) Method. Methods, 2001. 25(4): p. 402–8.

73. Sato, T., et al., Comparable Initial Engagement of Intracellular Signaling Pathways by Parathyroid Hormone Receptor Ligands Teriparatide, Abaloparatide, and Long-Acting PTH. JBMR Plus, 2021. 5(5): p. e10441.

74. Nagai, S., et al., Acute down-regulation of sodium-dependent phosphate transporter NPT2a involves predominantly the cAMP/PKA pathway as revealed by signaling-selective parathyroid hormone analogs. J Biol Chem, 2011. 286(2): p. 1618–26.

