## Supplemental Table 2 for "Regulation of intracellular cAMP levels in osteocytes by mechano-sensitive focal adhesion kinase via PDE8A"

| | | | |
|---|---|---|---|
| | Compound IC50 (M) | Control Compound IC50 (M) | Control compound ID |
| Target: | VS-6063 | | |
| PDE1A | No inhibition | 1.24E-05 | IBMX |
| PDE1B | No inhibition | 2.73E-04 | IBMX |
| PDE1C | No inhibition | 2.36E-04 | IBMX |
| PDE2A | 3.80E-06 | 6.03E-06 | IBMX |
| PDE3A | No inhibition | 8.83E-06 | IBMX |
| PDE3B | No inhibition | 1.54E-05 | IBMX |
| PDE4A | No inhibition | 5.45E-05 | IBMX |
| PDE4B | >1.00E-05 | 1.60E-05 | IBMX |
| PDE4C | No inhibition | 6.06E-05 | IBMX |
| PDE4D | No inhibition | 4.45E-05 | IBMX |
| PDE7A | No inhibition | 5.47E-06 | BRL-50481 |
| PDE8A | No inhibition | 3.33E-05 | TC-E 5005 |
| PDE10A | No inhibition | 7.72E-08 | TC-E 5005 |
| | No inhibition | | |

* Empty cells indicate no inhibition or compound activity that could not be fit to an IC50 curve.
