## Supplemental Table 3 for "Regulation of intracellular cAMP levels in osteocytes by mechano-sensitive focal adhesion kinase via PDE8A"

| qRT-PCR primers | Forward |
|---|---|
| Sost | GCCTCATCTGCCTACTTGTG |
| β-actin | CCTCTATGCCAACACAGTGC |

| CRISPR guide | Forward |
|---|---|
| FAK-KO1 (human) | CACCGATGTGGGAGATACTGATGCA |
| FAK-KO2 (human) | CACCGTCTGATGATAAATGACTGCG |
| FAK-KO3 (human) | CACCGCGATCATACTGGGAGATGCG |
| FAK-KO (mouse) | CACCGCAAGCAGCCTACTTACTCGA |
| PDE8A- KO (human) | CACCGCTGTGGTGAATATAACTCAG |

| FAK- shRNAs | |
|---|---|
| 1 | CCGATTGGAAACCAACATATA |
| 2 | GATGTTGGTTTAAAGCGATTT |

| Reverse |
|---|
| CTGTGGCATCATTCCTGAAG |
| ACATCTGCTGGAAGGTGGAC |

| Reverse |
|---|
| AAACTGCATCAGTATCTCCCACATC |
| AAACCGCAGTCATTTATCATCAGAC |
| AAACCGCATCTCCCAGTATGATCGC |
| AAACTCGAGTAAGTAGGCTGCTTGC |
| AAACCTGAGTTATATTCACCACAGC |
